# Human endogenous retrovirus HERV-K(HML-2) RNA causes neurodegeneration through Toll-like receptors

**DOI:** 10.1101/721241

**Authors:** Paul Dembny, Andrew G. Newman, Manvendra Singh, Michael Hinz, Michal Szczepek, Christina Krüger, Robert Adalbert, Omar al-Dzaye, Thorsten Trimbuch, Thomas Wallach, Gunnar Kleinau, Katja Derkow, Bernhard C. Richard, Carola Schipke, Claus Scheidereit, Douglas Golenbock, Oliver Peters, Michael Coleman, Frank L. Heppner, Patrick Scheerer, Victor Tarabykin, Klemens Ruprecht, Zsuzsanna Izsvák, Jens Mayer, Seija Lehnardt

## Abstract

Although human endogenous retroviruses (HERVs) represent a substantial proportion of the human genome and some HERVs have been suggested to be involved in neurological disorders, little is known about their biological function and pathophysiological relevance. HERV-K(HML-2) comprises evolutionarily young proviruses transcribed in the brain. We report that RNA derived from an HERV-K(HML-2) *env* gene region binds to the human RNA-sensing Toll-like receptor (TLR) 8, activates human TLR8, as well as murine Tlr7, and causes neurodegeneration through TLR8 and Tlr7 in neurons and microglia. HERV-K(HML-2) RNA introduced extracellularly into the cerebrospinal fluid (CSF) of either C57BL/6 wild-type mice or *APPPS1* mice, a mouse model for Alzheimer’s disease (AD), resulted in neurodegeneration. Tlr7-deficient mice were protected against neurodegenerative effects, but were re-sensitized towards HERV-K(HML-2) RNA when neurons ectopically expressed murine Tlr7 or human TLR8. Accordingly, transcriptome datasets of human brain samples from AD patients revealed a specific correlation of upregulated HERV-K(HML-2) and TLR8 RNA expression. HERV-K(HML-2) RNA was detectable more frequently in CSF from AD individuals compared to controls. Our data establish HERV-K(HML-2) RNA as an endogenous ligand for human TLR8 and murine Tlr7 and imply a functional contribution of specific human endogenous retroviral transcripts to neurodegenerative processes such as AD.

## Introduction

Neurodegenerative diseases are characterized by progressive loss of neurons, but the mechanisms underlying the deleterious spread of neuronal injury remain unclear. Toll-like receptors (TLRs) play a crucial role in regulating immunity against pathogens. Nevertheless, several studies also suggest a role for TLRs in non-infectious CNS inflammation and injury, in which pathogen-associated molecules are not detectable. Indeed, injured tissue and dying cells release host-derived molecules such as components of the extracellular matrix, heat shock proteins and RNA, which can act as endogenous ligands for TLRs (1). Contribution of TLRs to CNS damage has been reported in several mouse models of neurological disorders including Alzheimer’s disease (AD), amyotrophic lateral sclerosis (ALS), and multiple sclerosis (MS) (2). However, the endogenous ligands involved in TLR activation, the associated signaling pathways, and the cellular mechanisms by means of which TLRs lead to tissue injury in these different pathological contexts remain elusive. Based on our previous reports that TLR signaling can result in CNS injury (3–5), we postulate that processes triggered by endogenous ligands, which are potentially derived from injured CNS cells, may contribute to further neuronal injury in various forms of (inflammatory) neurodegenerative diseases, independently of the initial cause of the respective disease.

TLRs are membrane-bound receptors composed of an ectodomain constituted by leucin-rich repeats, which is involved in ligand binding and receptor activation, a transmembrane domain, and a cytoplasmic Toll/IL-1 receptor (TIR) domain. The TIR domain interacts with adaptor molecules, such as MyD88, which couple to downstream protein kinases and ultimately lead to the activation of transcription factors such as NF-κB, MAPKs, and IFN-regulatory factor family members. The latter in turn induce genes involved in inflammatory responses. The TLR family consists of 10 members in humans and 12 members in mice, of which TLRs 1-9 are conserved in their amino acid sequences in both species (6).

The genomes of vertebrates contain endogenous retroviruses, which are remnants of ancestral germline infections by exogenous retroviruses (7). Although human endogenous retroviruses (HERVs) comprise a substantial proportion of the human genome and some HERVs have been suggested to be involved in inflammatory and neurodegenerative processes, their biological functions and pathophysiological relevance remain largely unexplored. The multi-copy HERV-K(HML-2) group comprises numerous human-specific proviruses that are transcriptionally active in the brain (8–10). Although HERV-K(HML-2) encodes various former retroviral proteins and even retrovirus-like particles, no replication-competent HML-2 alleles have been identified as yet (11, 12). While HERV-K(HML-2) RNA is detected in various human tissue and cell types, transcription patterns of specific HERV-K(HML-2) proviruses differ considerably between tissue and cell types and also inter-individually, the latter due to presence/absence of alleles of HERV-K(HML-2) elements (9, 13–18). Deregulated HERV-K(HML-2) transcription has been reported for various diseases, including inflammatory and neurological disorders (19–21). HERV-derived RNA is present in blood, cerebrospinal fluid (CSF), and brain samples of patients with CNS diseases, such as MS, ALS, and Creutzfeldt-Jacob disease (21–24). Nonetheless, a direct signaling role of HERV RNA in cellular activation has not been demonstrated to date. Subsets of HERV-K(HML-2) proviruses encode, on the RNA level, the sequence 5’-GUUGUGU-3’ within the *envelope* (*env*) gene region while an evolutionarily younger HERV-K(HML-2) subset harbors a mutated sequence (5’-GUUG*C*GU-3’) in that position (see below). Notably, these motifs resemble the core of the GU-rich sequence that is responsible for species-specific recognition via Tlr7 (mice) and TLR8 (human) (25, 26).

We report here that extracellular HERV-K(HML-2) RNA is a potent activator of human TLR8 (hTLR8) and murine Tlr7 (mTlr7). While HERV-K(HML-2) RNA induces an inflammatory response in microglia and macrophages, stimulation of neuronal hTLR8 and mTlr7 leads to apoptosis in these cells. Injection of HERV-K(HML-2) transcripts into the CSF of mice induces neurodegeneration in an hTLR8- and mTlr7-dependent fashion. Correspondingly, we observed a higher frequency of HERV-K(HML-2) RNA presence in CSF and a parallel upregulation of HERV-K(HML-2) and TLR8 RNA expression in brains of patients with AD. Thus, our data implicate a contribution of HERV-K(HML-2) transcripts and TLR signaling to neurodegenerative processes such as AD.

## Results

### Extracellular HERV-K(HML-2) RNA activates Tlr7 and TLR8

TLR7 and TLR8 respond primarily to GU- or U-rich single-stranded RNA (ssRNA) and short interfering RNAs (25, 27–29). Among the known RNA ligands for TLR7 and TLR8 is the 20-nucleotide ssRNA40 derived from the exogenous retrovirus HIV (25, 26). We found that a subset of HERV-K(HML-2) proviruses (abbreviated as HERV-K in the following) contains sequences similar to ssRNA40, specifically a GUUGUGU motif within the *env* gene region (Supplementary Fig. 1) that is present in the core of ssRNA40 and is responsible for TLR7 and TLR8 activation (25, 26). Thus, we postulated that HERV-K RNA acts as an endogenous signaling activator of TLR7 and TLR8. To test this hypothesis, in a first step, we investigated the response of mTlr7-expressing microglia and macrophages (25, 30) to HERV-K RNA, using a synthetic 22-nucleotide containing the GUUGUGU motif (referred to as HERV-K) matching the *env* region of HERV-K. Following incubation with HERV-K both murine microglia (Fig. 1a) and bone marrow-derived macrophages (BMDM, Supplementary Fig. 2a) released cytokines and chemokines such as TNF-α (Fig. 1a, Supplementary Fig. 2a) and CXCL1 (Supplementary Fig. 2b) in a dose- and time-dependent fashion, and this response required the expression of mTlr7. In microglia, this response also required the intracellular adaptor protein Myd88 (Fig. 1a, Supplementary Fig. 2b). The HERV-K RNA effect was dependent on the GU-rich core, as a control oligoribonucleotide matching a sequence located upstream of the GUUGUGU motif within the *env* region of HERV-K (referred to as HERV-K (-GU) in the following) did not induce TNF-α and CXCL1 in microglia and macrophages (Fig. 1a, Supplementary Fig. 2a, b). The TLR ligands lipopolysaccharide (LPS, Tlr4), loxoribine (Tlr7) and poly(I:C) (Tlr3) served as positive controls for TLR-mediated cytokine and chemokine induction. The response of microglia deficient for both Tlr2 and Tlr4 was similar to that of wild-type cells after exposure to HERV-K RNA, excluding the possibility of contamination of the synthetic HERV-K oligoribonucleotide with LPS or Tlr2 ligands (Supplementary Fig. 2c). Human macrophages differentiated from THP-1 cells also responded to HERV-K RNA by TNF-α release in a sequence-, dose- and time-dependent manner (Fig. 1b). To test whether the canonical TLR-NF-κB pathway is involved in HERV-K RNA-induced signaling, we analyzed murine microglia and BMDM treated with HERV-K RNA by electrophoretic mobility shift assay (Fig. 1c, Supplementary Fig. 2d). HERV-K RNA potently induced NF-κB in microglia and BMDM, comparable to the positive control LPS and dependent on Tlr7 expression (Fig. 1c, Supplementary Fig. 2d), suggesting that HERV-K RNA directly activates Tlr7. Likewise, human macrophages responded to extracellular HERV-K RNA by NF-κB activation, although to a much lesser extent than to the one induced by LPS (Fig. 1d). Specificity of HERV-K RNA-induced NF-κB activation in microglia and macrophages was confirmed by the detection of the supershifted transcription factor subunits p50 and p65 and of IκB kinase phosphorylation after incubation with HERV-K RNA by western blot (Supplementary Fig. 2e, f).

**Fig. 1.**
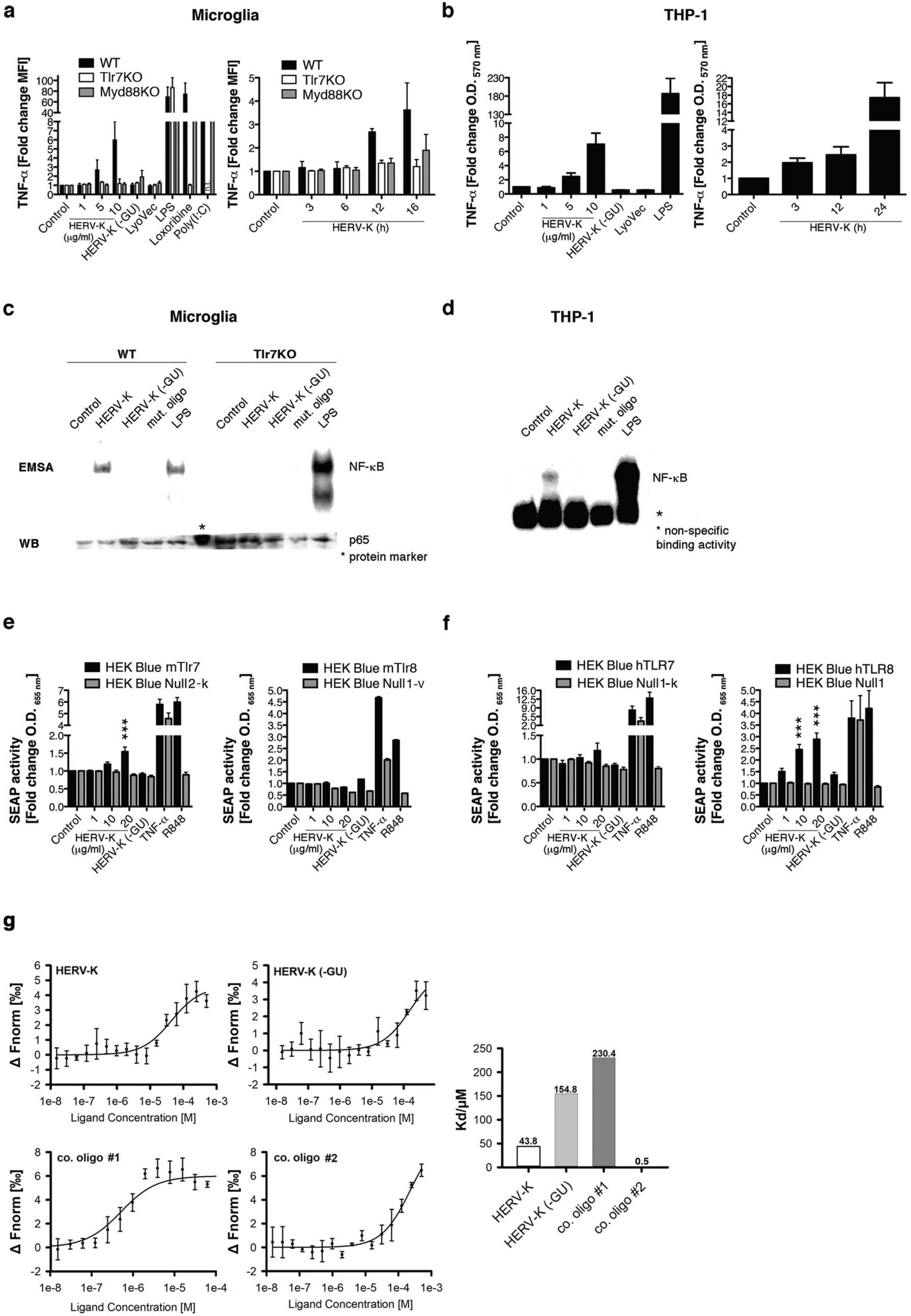
HERV-K(HML-2)-derived oligoribonucleotides bind to TLR8 and induce TNF-α release from microglia and macrophages via Tlr7 and TLR8. (**a**) Microglia from C57BL/6 (wild-type, WT), Tlr7KO or Myd88KO mice and (b) macrophages differentiated from THP-1 cells were incubated for 12 h with various doses of HERV-K(HML-2) (referred to as HERV-K) oligoribonucleotide containing a GUUGUGU motif present in the *env* region (HERV-K, left) or with 5 µg/ml of HERV-K for various durations (right). Untreated cells (control) and an oligoribonucleotide with a sequence located upstream from the GU-rich motif within the HERV-K *env* region (HERV-K (-GU), 10 µg/ml) served as blank and negative control, respectively. Incubation with the transfection agent LyoVec alone served as a further negative control. LPS (100 ng/ml), loxoribine (1 mM), and poly(I:C) (100 ng/ml) were used as established ligands for TLR4, TLR7, and TLR3 activation, respectively. Subsequently, TNF-α amounts in the culture supernatants were determined by immuno multiplex assay. Data are pooled from three experiments. n.t. not tested. (**c**) Purified microglia and (**d**) macrophages differentiated from THP1-cells were incubated for 2 h with 5 µg/ml HERV-K, 5 µg/ml HERV-K (-GU), or a mutant oligoribonucleotide, in which the GU content had been reduced. 1 µg/ml LPS served as a positive control. Subsequently, protein lysates were assayed for NF-κB activation by EMSA. Western blot using an antibody against p65 confirmed equal loading of probes. One representative experiment of at least three independent experiments is shown. HEK-Blue cells stably co-expressing murine (**e**) or human (**f**) TLR7 or TLR8 and an NF-κB/AP1-inducible secreted embryonic alkaline phosphatase (SEAP) reporter gene were incubated for 48 h with various doses of HERV-K, as indicated. HERV-K (-GU) (20 µg/ml) served as a negative control. R848 served as a ligand control for TLR7/TLR8 activation, while TNF-α served as a positive control for SEAP induction. HEK-Blue cells lacking TLR7/TLR8 expression (*Null* cells) were tested as negative control cell line. Data are pooled from three to seven experiments. Results are presented as mean ± s.e.m. \*\*\**P* < 0.001 over HERV-K dose, as indicated, compared to control (one-way ANOVA with Bonferroni’s post hoc analysis). (**g**) Binding affinity measurements of the purified polyhistidine-tagged TLR8 protein and synthetic oligoribonucleotide variants using microscale thermophoresis (MST). TLR8-RNA interaction was monitored by titrating RNA from 500 µM to 30 nM (HERV-K, HERV-K (-GU), control oligoribonucleotide #1) and 62.5 µM to 3.8 nM (control oligoribonucleotide #2) against 50 nM RED-tris-NTA-labeled TLR8 measured with the Nanotemper Monolith NT.115 device (left). *K_d_* values were calculated from dose response curves, which were calculated from titration experiments (right). Results are presented as mean ± s.d. *n* = 3.

To systematically validate the ability of HERV-K RNA to activate the TLR signaling pathway in both murine and human cell systems and to distinguish between the responses mediated by TLR7 and/or the phylogenetically and structurally highly related TLR8 (25) we employed HEK 293 cells, stably co-expressing murine (Fig. 1e) or human (Fig. 1f) TLR7 or TLR8 and an NF-κB/AP1-inducible secreted embryonic alkaline phosphatase reporter gene. As expected, HERV-K RNA induced NF-κB in HEK 293 cells expressing mTlr7 in a sequence-dependent manner. In contrast, HEK 293 cells expressing hTLR7 did not respond to HERV-K RNA treatment. Instead, HERV-K RNA potently induced NF-κB in HEK 293 cells expressing hTLR8 in a sequence-specific fashion, while cells expressing mTlr8 did not respond to HERV-K RNA.

To determine whether HERV-K RNA serves as an immediate ligand for hTLR8, we made use of microscale thermophoresis (Fig. 1g). Interaction between HERV-K oligoribonuleotide with hTLR8 protein yielded a dissociation constant (*K_d_*) of 43.8, confirming direct binding of HERV-K RNA harboring the GU-rich motif to hTLR8. Comparatively, interaction between HERV-K (-GU) RNA and hTLR8 resulted in a *K_d_* of 154.8, while sequence-modified control oligoribonucleotides bound to the receptor with a *K_d_* of 230.4 and a *K_d_* of 0.5, respectively.

Taken together, HERV-K RNA serves as a ligand for Tlr7 and TLR8, in mice and humans, respectively, thereby inducing the canonical TLR pathway involving Myd88 and NF-κB in microglia and macrophages.

### HERV-K RNA causes neuronal injury via TLR signaling *in vitro*

TLR7 and TLR8 are primarily expressed in immune cells (25, 27), but were also detected in murine neurons (5, 31, 32). We tested whether these receptors are expressed in human neurons. As is the case for murine neurons, both TLR7 and TLR8 were readily detectable in neurons derived from the human neuroblastoma cell line SH-SY5Y by FACS analysis (Fig. 2a). We observed that biotinylated HERV-K oligoribonucleotide enters neurons without transfection (Fig. 2b). To investigate whether extracellular HERV-K RNA might affect primary neurons, cortical neurons from wild-type and Tlr7-deficient mice were purified and incubated with HERV-K oligoribonucleotides containing or lacking the GUUGUGU motif (HERV-K or HERV-K (-GU)). HERV-K RNA induced injury and loss in wild-type neuronal cultures, as determined by immunostaining with antibodies against dendrites and neuronal nuclei (Fig. 2c). This effect was dose- and time-dependent, exceeding the activity of the established Tlr7 ligand loxoribine (Fig. 2d). HERV-K RNA-induced neurotoxicity was sequence-specific, since deletion of the GU motif in the control oligoribonucleotide eliminated cell death induction (Fig. 2d). It followed that if HERV-K RNA interacts with Tlr7, Tlr7 mutant mice would be resistant to extracellular HERV-K RNA and indeed, whereas HERV-K RNA induced cell death in wild-type neurons, Tlr7-deficient neurons were unaffected by HERV-K RNA treatment (Fig. 2c, d). To determine the molecular process of the observed neurotoxicity, neurons were analyzed for the expression of active caspase-3, a downstream effector of apoptosis. HERV-K RNA treatment caused an increase in the expression of active caspase-3 in wild-type cultures but not in Tlr7-deficient neurons (Fig. 2e). Next, we asked, which downstream TLR signaling module conducts HERV-K RNA-induced neurotoxicity. Since the canonical Tlr7-mediated signaling pathway involves Myd88, we analyzed the response of Myd88 mutant cell cultures to treatment with HERV-K RNA (Fig. 2d). Similar to wild-type neurons Myd88-deficient neurons underwent dose- and time-dependent cell death in response to HERV-K RNA, indicating that Myd88 is not required for neuronal injury induced by HERV-K oligoribonucleotide. Correspondingly, HERV-K RNA did not induce canonical activation of NF-κB in neurons (Supplementary Fig. 3a, b).

**Fig. 2.**
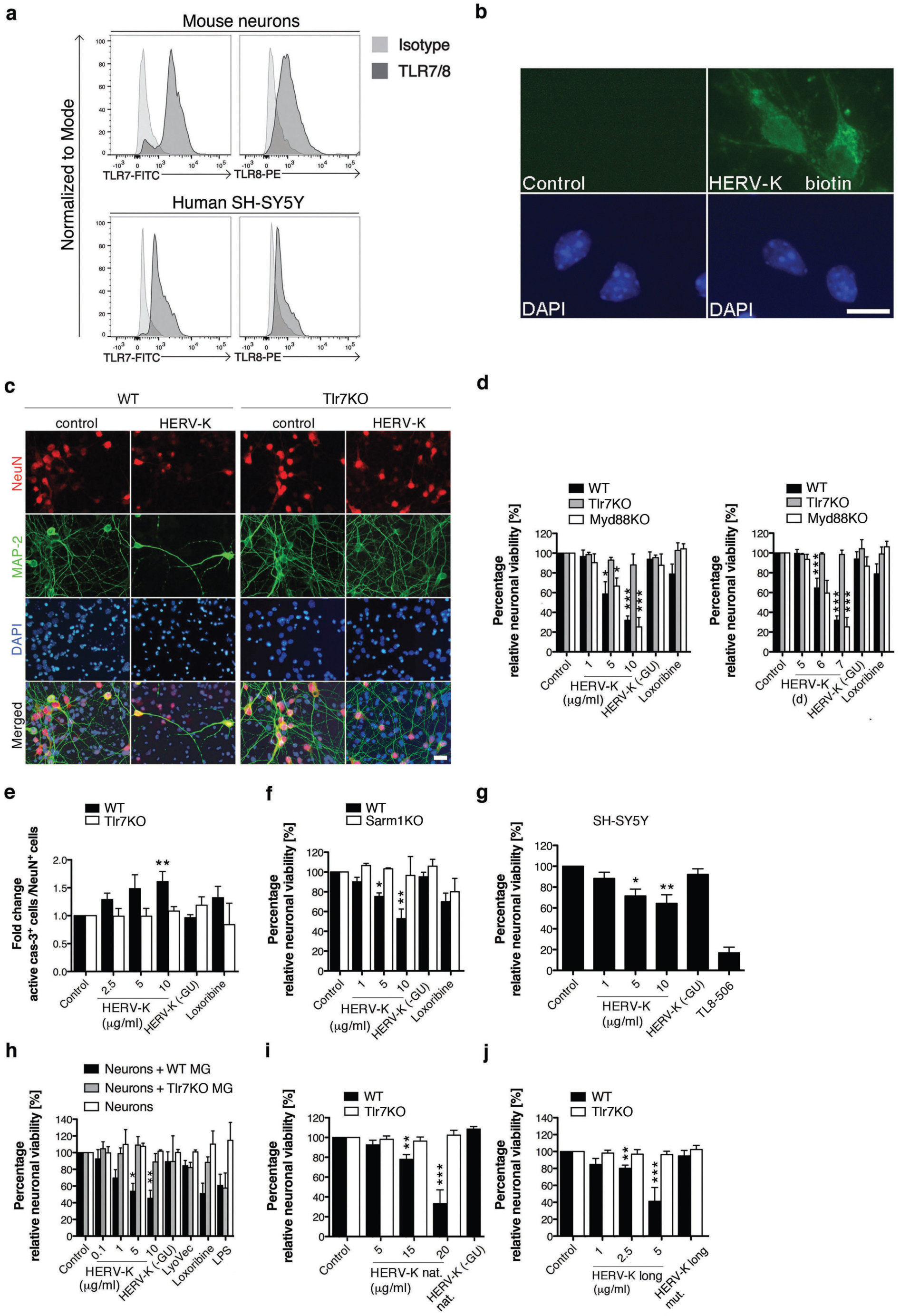
Extracellular HERV-K RNA induces neuronal cell death through Tlr7, Sarm1 and caspase-3. (**a**) Murine cortical neurons (upper panel) and human SH-SY5Y cells (lower panel) were stained with anti-TLR7-FITC (left) or anti-TLR8-PE (right) antibodies and analyzed by flow cytometry. The respective isotype was used as a negative control. (**b**) Murine neurons were incubated with 5 µg/ml biotinylated HERV-K oligoribonucleotide harboring a GUUGUGU motif (HERV-K) or PBS (control) for 12 h and subsequently stained with Avidin-Alexa 488 and DAPI. Scale bar, 10 µm. (**c**) Wild-type (WT) and Tlr7KO neurons were incubated with 5 µg/ml HERV-K for 7 d. Untreated cells served as control. Cells were immunostained with NeuN and MAP-2 antibodies and DAPI. Scale bar, 10 µm. (**d**) WT, Tlr7KO and Myd88KO neurons were incubated with various doses of HERV-K for 7 d (left) or with 5 µg/ml HERV-K for various durations (right), and relative neuronal viability was assessed (*P* = 0.0001 and *P* < 0.0001 over dose and time, respectively, WT all groups; *P* < 0.0001 and *P* = 0.0006 over dose and time, respectively, Myd88KO all groups; not significant over Tlr7KO all groups, Kruskal-Wallis test). \**P* < 0.05, \*\*\**P* < 0.001 compared with control (*n* = 4-11). (**e**) WT and Tlr7KO neurons were incubated with various doses of HERV-K for 5 d and immunostained for active caspase-3, NeuN antibody and DAPI. Subsequently, caspase-3-positive cells were quantified (*P* = 0.0005 over WT all groups and not significant over Tlr7KO all groups, Kruskal-Wallis test). \*\**P* < 0.01 compared with control (*n* = 5-6). (**f**) WT and Sarm1KO neurons were incubated with different doses of HERV-K for 7 d, and relative neuronal viability was assessed (*P* = 0.0006 over WT all groups and not significant over Sarm1KO all groups, Kruskal-Wallis test). \**P* < 0.05, \*\**P* < 0.01 compared with control (*n* = 4-10). (**g**) SH-SY5Y cells were incubated with various doses of HERV-K for 3 d, and relative neuronal viability was assessed (*P* = 0.0015 over all groups, Kruskal-Wallis test). \**P* < 0.05, \*\**P* < 0.01 compared with control (*n* = 6). (**h**) Purified WT neurons and neuron/microglia (+ WT MG or + Tlr7KO MG) co-cultures were incubated with HERV-K for 3 d, and relative neuronal viability was assessed (*P* = 0.009 over neurons + WT MG all groups; not significant over neurons + Tlr7KO MG all groups; not significant over neurons all groups, Kruskal-Wallis test). \**P* < 0.05, \*\**P* < 0.01 compared with control (*n* = 5-6). (**i**) WT and Tlr7KO neurons were incubated with indicated doses of non-phosphorothioated HERV-K oligoribonucleotide for 7 d, and relative neuronal viability was assessed (*P* < 0.0001 over WT all groups and not significant over Tlr7KO all groups, Kruskal-Wallis test). \*\**P* < 0.01, \*\*\**P* < 0.001 compared with control (*n* = 3-12). (**j**) WT and Tlr7KO neurons were incubated with indicated doses of a 50-nucleotide HERV-K oligoribonucleotide (HERV-K long) or with a scrambled 50-nucleotide HERV-K (HERV-K long mut.) for 7 d, and relative neuronal viability was assessed (*P* < 0.0001 over WT all groups and not significant over Tlr7KO, Kruskal-Wallis test). \*\**P* < 0.01, \*\*\**P* < 0.001 compared with control (*n* = 4-11). In all panels, unless stated otherwise, 10 µg/ml HERV-K (-GU) oligoribonucleotide served as a negative control, whereas loxoribine served as an established TLR7 activator. Results are presented as mean ± s.e.m. *P* values for relevant groups, as indicated, were determined by Kruskal-Wallis test with Dunn’s post-hoc analysis.

Sterile alpha and TIR domain-containing 1 (SARM1) is a member of the Toll/ILR (TIR) domain-containing adaptor protein family and was recently identified as a mediator of axonal degeneration and neurodegeneration (33–35). To test the role of SARM1 as an adaptor protein in HERV-K RNA-triggered neuronal cell death, neurons from wild-type and Sarm1-deficient mice were incubated with HERV-K oligoribonucleotide. In contrast to wild-type neurons Sarm1-deficient neurons were resistant to HERV-K RNA (Fig. 2f), indicating that Sarm1 is a crucial mediator for neuronal injury induced by HERV-K/TLR signaling.

A relevant contamination of the murine neuronal cultures analyzed above with glial cells was ruled out by immunostaining (5). To confirm the cell-autonomous nature of neuronal cell death induced by HERV-K RNA and to test HERV-K RNA-induced toxic effects in human-derived neurons, SH-SY5Y cells were incubated with HERV-K oligoribonucleotide. SH-SY5Y cells incubated with HERV-K RNA underwent sequence-specific cell death similar to that observed in primary murine neuronal cell cultures (Fig. 2g). Likewise, HERV-K RNA did not induce NF-κB activation in SH-SY5Y cells (Supplementary Fig. 3c). Thus, using highly purified neurons and the neuronal cell line SH-SY5Y, our data indicate that microglia are not required for HERV-K RNA-induced neuronal injury, despite the ability of HERV-K RNA to induce an inflammatory response in microglia (see Fig. 1a, Supplementary Fig. 2b). Nevertheless, as microglia can mediate neuronal injury through TLRs (3), we compared HERV-K RNA-mediated effects in cultures of purified neurons with co-cultures of neurons and microglia derived from wild-type or Tlr7-deficient mice (Fig. 2h) after 3 days of incubation, when HERV-K RNA-induced loss of neurons was not observed in the experiments described above (see Fig. 2d for comparison). In the presence of wild-type microglia, HERV-K RNA-induced neuronal cell death occurred three days earlier and at lower HERV-K doses compared to the toxic effects observed in purified neurons, pointing to an additional impact on neuronal survival mediated by an inflammatory environment. This microglia-enhanced neuronal cell death required the expression of Tlr7 in microglia, as expected (Fig. 2h).

We used HERV-K oligoribonucleotide stabilized by phosphorothioate linkages throughout the study, but confirmed that unmodified HERV-K RNA also induced neurotoxicity dependent on Tlr7 *in vitro* (Fig. 2i). The specificity of HERV-K RNA as the causative agent of the observed neurotoxic effects was additionally confirmed by using a larger HERV-K oligoribonucleotide (50-nucleotide, referred to as HERV-K long, Fig. 2j), which induced sequence-specific and Tlr7-dependent cell death in neuronal cultures. These neurotoxic effects seemed to be enhanced compared to those observed in the toxicity assays using the 22-oligoribonucleotide (see Fig. 2d for comparison).

Taken together, HERV-K RNA induces neuronal apoptosis via TLR and SARM-1 signaling in these cells. This cell-autonomous neurotoxic effect can be enhanced by an inflammatory response mediated by microglia in response to HERV-K RNA.

### Endogenous HERV-K transcripts are released from injured neurons and induce further neurodegeneration *in vitro*

Having shown that oligonucleotides specifically derived from the HERV-K *env* region trigger neurotoxicity, we sought to determine whether endogenous full-length HERV-K transcripts are capable of inducing neuronal injury. Analysis of SH-SY5Y cells by RT-PCR confirmed that human-derived neurons harbor HERV-K transcripts containing the sequence GUUGUGU (Fig. 3a). The human teratocarcinoma cell line Tera-1, which transcribes HERV-K at a high level and produces HERV-K-encoded retroviral particles containing full-length HERV-K transcripts, some of which harbor GU-rich sequences including the motif GUUGUGU (12, 14), served as a positive control. Retroviral particles released from Tera-1 cells were isolated, and presence of HERV-K transcripts in retroviral particles was confirmed by endpoint RT-PCR (Fig. 3b). Subsequently, murine cortical neurons were incubated with virions containing HERV-K transcripts, which led to neuronal cell death that was dependent on Tlr7 expression in neurons (Fig. 3c). We then tested whether this virion-induced loss of neurons can be prevented by using an antagonist directed against the HERV-K *env* region containing the GUUGUGU motif (named HERV-K inhibitor, Fig. 3d). Whereas neurons without inhibitor treatment underwent cell death in response to virions, neurons were protected against cell death induced by the virion particles in the presence of the HERV-K inhibitor. In contrast, a negative control inhibitor consisting of a scrambled RNA sequence failed to provide such a protective effect (Fig. 3d). Thus, neuronal cell death mediated by Tera-1 cell-derived retroviral particles was specifically induced by endogenous HERV-K transcripts containing the GUUGUGU motif.

**Fig. 3.**
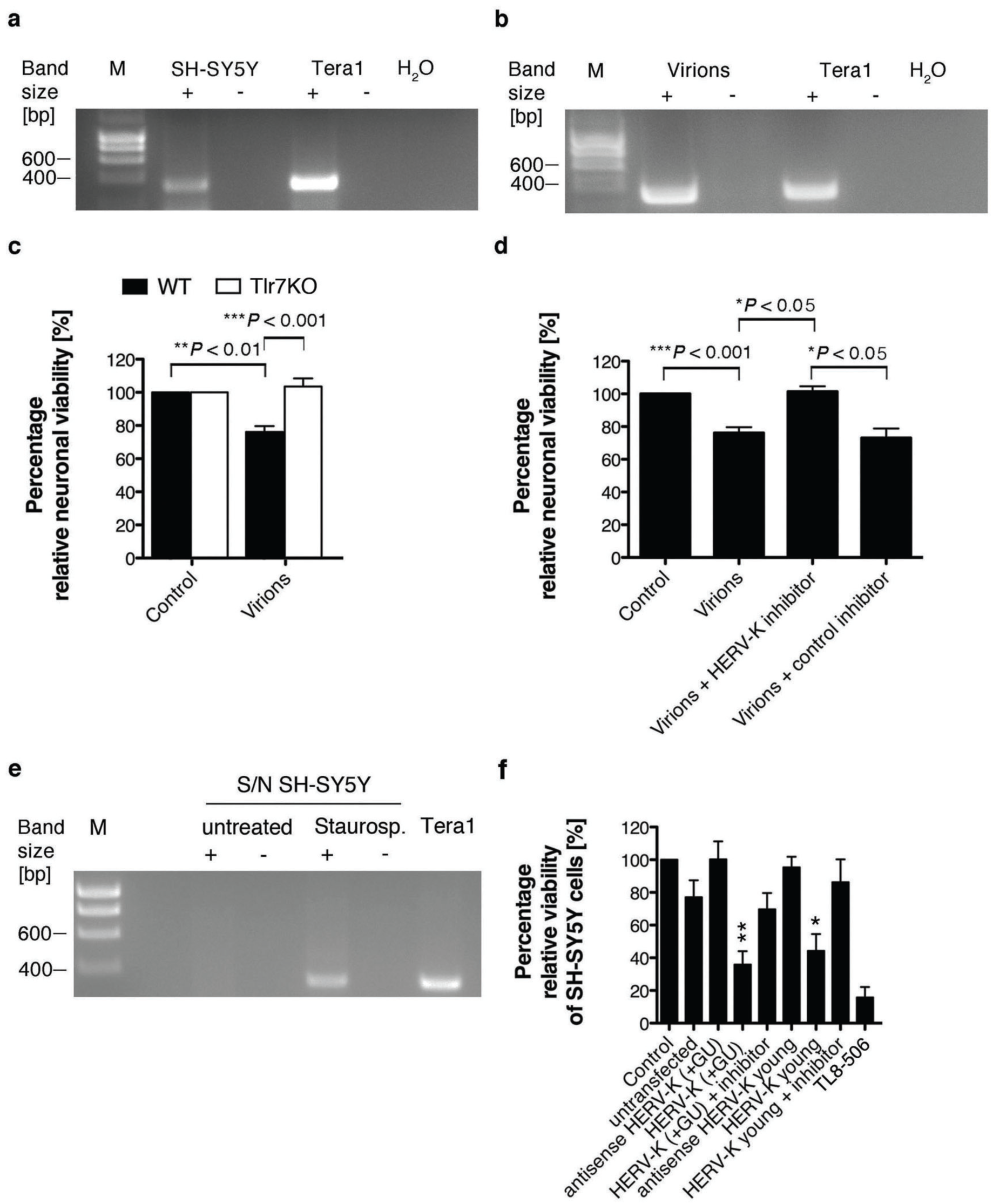
Endogenous HERV-K transcripts induce neuronal injury *in vitro*. (**a**) SH-SY5Y cells and (**b**) virion particles isolated from Tera-1 cells were analyzed by RT-PCR using primers amplifying a HERV-K *env* region harboring the motif GUUGUGU. Tera-1 cells served as positive control. (**c**) Neurons isolated from wild-type (WT) and Tlr7KO mice were incubated with 10 µl suspension of virions isolated from Tera-1 cells or were left untreated (control) for 6 d, and relative neuronal viability was determined (*P* < 0.0001 over all groups, Kruskal-Wallis test) (*n* = 6-10). (**d**) WT neurons were incubated with 10 µl suspension of virions isolated from Tera-1 cells for 6 d, with or without HERV-K inhibitor. A non-specific inhibitor served as control for inhibitor specificity. Subsequently, relative neuronal viability was assessed (*P* < 0.0001 over all groups, Kruskal-Wallis test) (*n* = 5-10). (**e**) Supernatants (S/N) from untreated and apoptotic neurons treated with staurosporine were analyzed by RT-PCR using primers specific for HERV-K. (**f**) 20 µg/ml RNA isolated from untransfected SH-SY5Y cells (control), from SH-SY5Y cells overexpressing HERV-K RNA harboring a GUUGUGU motif (HERV-K (+ GU)), a GUUG*C*GU motif (HERV-K young), or the respective antisense sequence were used for the incubation of freshly plated SH-SY5Y cells for 4 d, with or without HERV-K inhibitor. Untreated SH-SY5Y cells served as a further negative control, whereas TL8-506 served as a positive control for TLR8 activation. Relative neuronal viability was assessed (*P* = 0.0001 over all groups, Kruskal-Wallis test). Results are presented as mean ± s.e.m. \**P* < 0.05, \*\**P* < 0.01 compared with control (*n* = 5). *P* values, as indicated, were determined by the Kruskal-Wallis test with Dunn’s post hoc analysis.

Given that injured tissues release molecules into their local environment, thereby activating innate immune receptors (1, 4), we hypothesized that HERV-K RNA might be released during neuronal injury and subsequently act on neighboring neurons. In a chain reaction, these neurons could accelerate neuronal damage by releasing neurotoxic HERV-K RNA. To test whether injured human neurons release HERV-K RNA apoptosis was induced in SH-SY5Y cell cultures by adding staurosporine. Culture supernatants were freed of cell remnants by centrifugation and analyzed for the presence of HERV-K RNA by RT-PCR (Fig. 3e). Whereas HERV-K RNA was not detected in supernatants from untreated neurons, HERV-K RNA was observed in supernatants from apoptotic neurons. We sought to confirm that endogenous HERV-K transcripts derived from human-derived neurons are capable of inducing neuronal injury. To do so, total RNA from native SH-SY5Y cells and SH-SY5Y cells overexpressing a 785-nucleotide covering the HERV-K *env* region and containing the GUUGUGU motif (HERV-K (+GU)) or the sequence GUUG*C*GU (HERV-K young), the latter being present in an evolutionarily younger HERV-K subset (Supplementary Fig. 1b), was isolated and incubated with freshly plated SH-SY5Y cells (Fig. 3f). Similar to synthetic HERV-K oligoribonucleotides, RNA isolated from cells overexpressing HERV-K *env* transcripts induced neuronal cell death in SH-SY5Y cell cultures, comparable to the TLR8 agonist TL8-506. Neurotoxicity was, at least in part, specifically induced by HERV-K transcripts containing either the GUUGUGU or the GUUG*C*GU motif as these overexpressed transcripts exceeded neurotoxic effects induced by RNA derived from untransfected SH-SY5Y cells, and RNA isolated from SH-SY5Y cells overexpressing the respective RNA in antisense failed to induce comparable cell death (Fig. 3f). Furthermore, the presence of HERV-K inhibitor (see above) protected neurons against cell death induced by HERV-K transcripts containing either the GUUGUGU or the GUUG*C*GU motif (Fig. 3f).

Taken together, these data indicate that endogenous HERV-K transcripts, which are released from both murine and human injured neurons, initiate a chain reaction causing further neurodegeneration.

### HERV-K RNA in cerebrospinal fluid induces neurodegeneration via mTlr7 and hTLR8 *in vivo*

To evaluate the role of HERV-K RNA as an activator of Tlr7 in neurons *in vivo*, we injected mice intrathecally with HERV-K oligoribonucleotide containing the GUUGUGU motif (HERV-K (+GU)), a control oligoribonucleotide lacking this motif (HERV-K (-GU)), or a mutant oligoribonucleotide, in which the GU-content had been reduced. Immunohistochemical analysis of the cerebral cortex 3 days after injection revealed that HERV-K (+GU) RNA induced marked axonal injury (Fig. 4a) and neuronal loss (Fig. 4b) in wild-type mice, whereas the HERV-K (-GU) and mutant oligoribonucleotides affected neither axon integrity nor the numbers of neurons. In contrast, Tlr7-deficient mice were protected from the neurodegenerative effects of HERV-K (+GU) RNA (Fig. 4a, b). Immunohistochemical analysis using an antibody against active caspase-3 confirmed the induction of apoptosis exclusively in the cerebral cortex of HERV-K (+GU)-treated wild-type animals (Fig. 4c). HERV-K (+GU) RNA-induced neurodegeneration increased with time, with immunohistochemistry revealing a loss of 22.4% of neurons in the cerebral cortex after 3 days (Fig. 4b) and 31.9% of neurons after 2 weeks (Fig. 4d, e), whereas Tlr7-deficient mice displayed no neuronal damage or loss during the whole observation period. Similarly, intrathecal injection of an HERV-K oligoribonucleotide containing the evolutionarily younger motif GUUG*C*GU (HERV-K young) also resulted in loss of cortical neurons dependent on mTlr7 (Supplementary Fig. 4). To confirm the specificity of HERV-K RNA as the causative factor, we injected wild-type mice intrathecally with the HERV-K inhibitor or a non-specific control inhibitor 16 hours before intrathecal injection of HERV-K (+GU) RNA or the mutant oligoribonucleotide. Immunohistochemical analysis of the cerebral cortex after 3 days revealed that pre-treatment with the HERV-K inhibitor abolished neurodegenerative effects induced by exogenous HERV-K (+GU) RNA (Fig. 4f). In contrast, pre-treatment with the non-specific control inhibitor did not protect neurons.

**Fig. 4.**
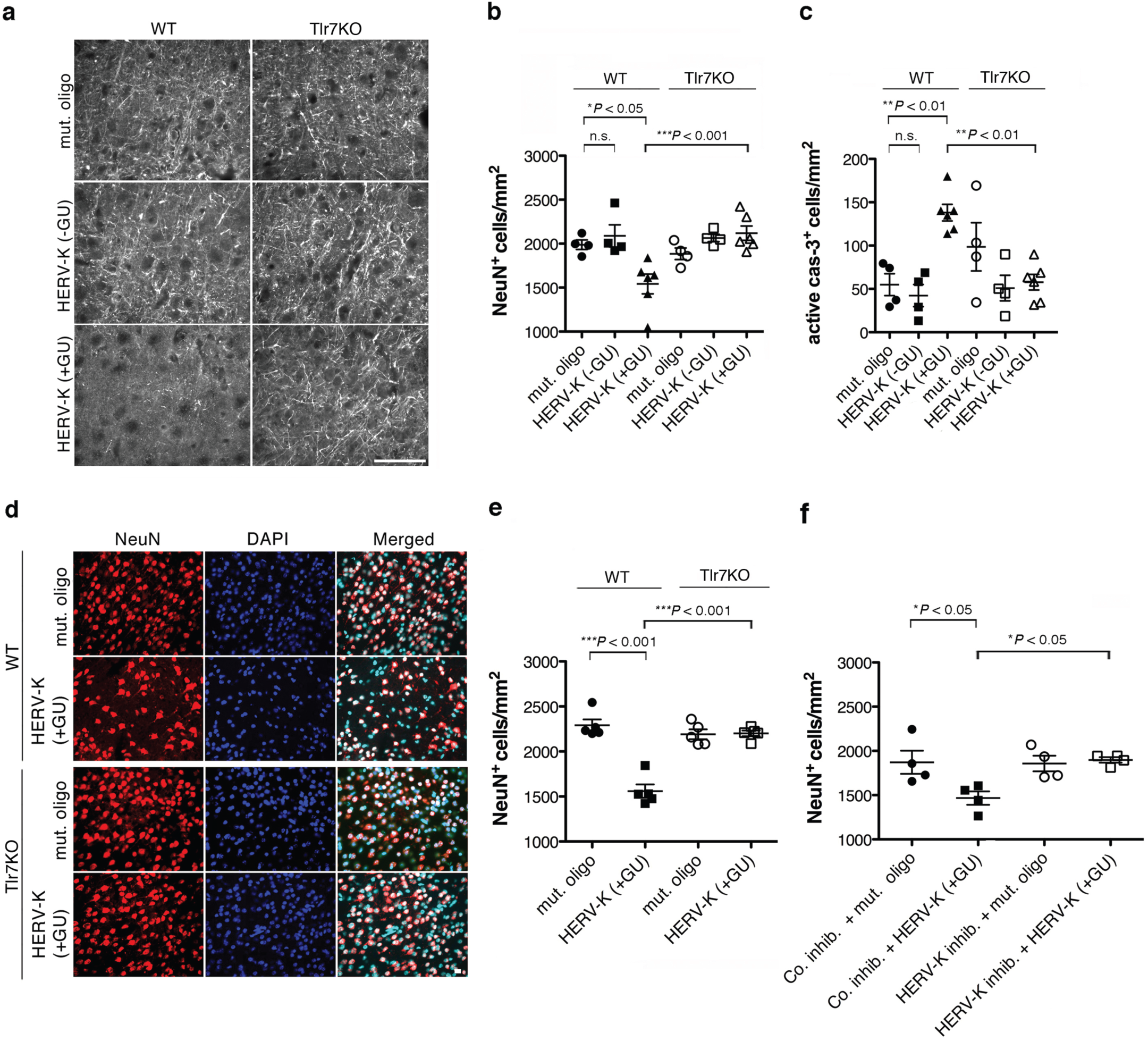
Intrathecal administration of HERV-K RNA causes neurodegeneration through Tlr7. 10 µg HERV-K, 10 µg HERV-K (-GU) oligoribonucleotide, or 10 µg mutant oligoribonucleotide were injected intrathecally into C57BL/6 (WT, HERV-K, *n* = 5-6; HERV-K (-GU), *n* = 4; mutant oligoribonucleotide, *n* = 4-5) or Tlr7-deficient (Tlr7KO, HERV-K, *n* = 5-6; HERV-K (-GU), *n* = 4; mutant oligoribonucleotide, *n* = 4-5) mice. (**a**-**c**) After 3 d, brain sections were immunostained with Neurofilament antibody (**a**, scale bar, 50 µm), NeuN antibody (**b**), or an antibody against active caspase-3 (**c**) and DAPI. NeuN-positive (**b**) and caspase-3-positive (**c**) cortical cells were quantified. (**d**, **e**) After 14 d, brain sections were immunostained with NeuN antibody and DAPI (**d**, scale bar, 10 µm), and NeuN-positive cortical cells were quantified (**e**). (**f**) 125 pmol of HERV-K inhibitor or nonspecific inhibitor (co. inhib.) were injected intrathecally into WT mice. After 16 h, mice were injected intrathecally with 10 µg of HERV-K or 10 µg of mutant oligoribonucleotide (for HERV-K inhibitor: HERV-K, *n* = 4; mut. oligo, *n* = 4; for co inhib: HERV-K, *n* = 4; mut. oligo, *n* = 4). After further 3 d, brain sections were immunostained with antibody to NeuN, and NeuN-positive cortical cells were quantified. Results are presented as mean ± s.e.m. A one-way ANOVA test yielded values of *P* = 0.0007 (**b**), *P* = 0.0003 (**c**), *P* < 0.0001 (**e**), and *P* = 0.0145 (**f**) over all groups. *P* values for relevant groups were determined by one-way ANOVA with Bonferroni’s post hoc test, as indicated. n.s., not significant.

To analyze whether extracellular HERV-K contributes to accumulation of microglia and Aβ plaque pathology - besides neuronal loss and neurofibrillary tangles two major hallmarks of Alzheimer’s disease (AD) (36, 37) - we injected amyloid-prone *APPPS1* mice (38) intrathecally with HERV-K RNA combined with or without HERV-K inhibitor treatment. Morphometric analysis revealed no significant changes with regard to Aβ plaque load in mice injected with HERV-K compared to sham animals (Supplementary Figure 5a). However, we observed a 66.6% increase in the number of microglia at 120 d of age in *APPPS1* mice injected with HERV-K RNA. Also, HERV-K oligoribonucleotide induced a loss of 31.9% of neurons in the cerebral cortex of *APPPS1* mice, as already observed for C57BL/6 mice (see above). Pre-treatment with HERV-K inhibitor before HERV-K injection abolished both the increased number of microglia and neurodegenerative effects (Supplementary Figure 5b). We note in this context that studies on extracellular HERV-K in *APPPS1* mice have to be considered with caution as *APPPS1* mice only partially recapitulate human AD, those mice lack HERV-K, and HERV-K expression must be regarded in a human-specific context. Nevertheless, our data from *APPPS1* mice corroborate our findings on HERV-K-induced neurodegeneration and microglial activation *in vivo*.

To further link TLR expression to HERV-K RNA-induced neurodegeneration, we reintroduced mTlr7 into the cerebral cortex of Tlr7-deficient mice by *in utero* electroporation. Since we found that hTLR8 also responds to HERV-K RNA (see Fig. 1) and since TLRs and their associated signaling pathways are highly conserved in different species including humans and mice (6), we also transfected Tlr7-deficient mice with hTLR8 in this experimental approach. Tlr7-deficient mouse embryos were electroporated with a plasmid encoding GFP and mTlr7 or hTLR8 or with a control vector expressing RFP (tdTomato) only at embryonic day (E) 14.5. At postnatal day (P) 19, HERV-K oligoribonucleotide containing the GUUGUGU motif (HERV-K) or mutant oligoribonucleotide was administered intrathecally, and cortices were analyzed 3 days later (Fig. 5). Where mTlr7 or hTLR8 was reintroduced into HERV-K RNA-treated Tlr7KO cortices, we observed a 58.9% and 50.1% loss, respectively, of GFP-positive neurons compared to mice injected with the control mutant oligoribonucleotide (Fig. 5a, b). Likewise, reintroduction of mTlr7 and hTLR8 resulted in increased numbers of cells positive for active caspase-3 in the cerebral cortex of mice injected with HERV-K RNA compared to mice injected with the mutant oligoribonucleotide (Fig. 5c). Unelectroporated NeuN-positive neurons were not affected by HERV-K RNA treatment (Fig. 5b), and injection of HERV-K RNA had no effect on neurons or the numbers of caspase-3-positive cells transfected with the control vector (Fig. 5a-c). These results confirm that the neurodegenerative activity of intrathecal HERV-K RNA requires neuronal mTlr7 or hTLR8. Moreover, these data demonstrate a functional role for a human TLR, namely hTLR8, expressed in neurons in HERV-K RNA-induced neurodegeneration *in vivo*.

**Fig. 5.**
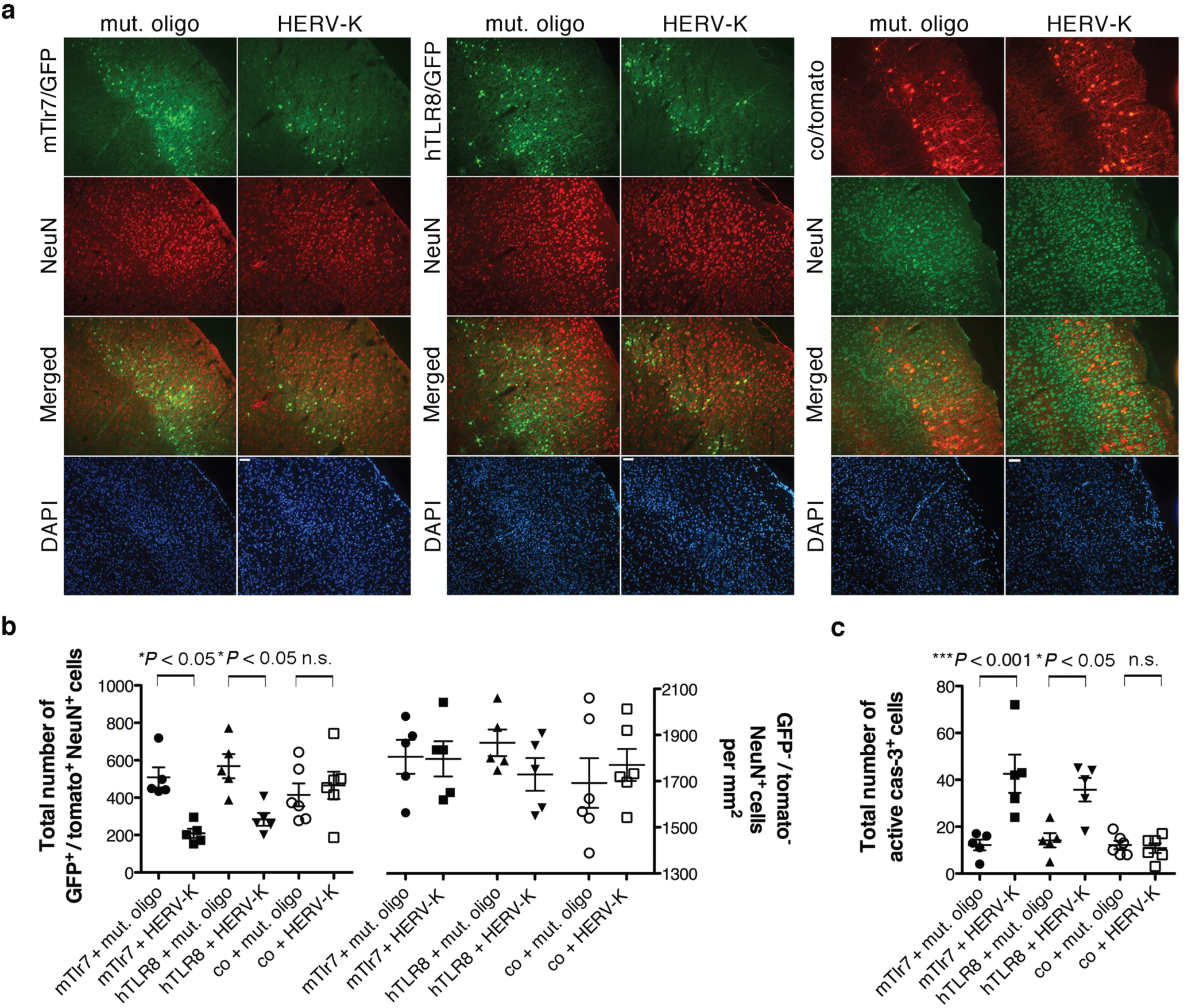
Transfection of murine Tlr7 or human TLR8 restores the neurodegenerative effect of HERV-K RNA *in vivo*. (**a**) Tlr7KO embryos (E14.5) were electroporated *in utero* with an expression vector for murine Tlr7 (Tlr7/GFP, left), an expression vector for human TLR8 (TLR8/GFP, middle) or control (co/RFP (td tomato), right). eGFP was coexpressed from an IRES cassette. At postnatal day 19, 10 µg of HERV-K RNA (Tlr7 vector, *n* = 5; TLR8 vector, *n* = 5; control vector, *n* = 6) or mutant oligoribonucleotide (Tlr7 vector, *n* = 5; TLR8 vector, *n* = 5; control vector, *n* = 6) were administered intrathecally and the cerebral cortex was analyzed by immunostaining with antibody to NeuN 3 d later. Scale bars represent 50 µm. (**b**) Quantitation of GFP^+^/tomato^+^ NeuN^+^ cells (left) and GFP^−^/tomato^−^ NeuN^+^ cells (right) in the electroporated cortex. (**c**) Sections of the electroporated cortex were immunostained with antibody to active caspase-3, and caspase-3^+^ cells were quantified. Five representative brain sections per mouse were analyzed. Results are presented as mean ± s.e.m. The one-way ANOVA test yielded values of (**b**) *P* = 0.0014 (left) and n.s. (right) and (**c**) *P* < 0.0001 over all groups. *P* values for relevant groups were determined by one-way ANOVA with Bonferroni’s post hoc test, as indicated. n.s., not significant.

### Correlation of upregulated HERV-K and TLR8 expression in Alzheimer’s disease

Differential expression of HERV-K in brains of individuals with neurodegenerative diseases including AD was previously observed (24, 39). To determine whether a correlation between HERV-K transcription and TLR7/8 expression in AD patients exists, we analyzed transcriptome datasets (40). The cohort, provided by the Mayo brain bank (https://www.synapse.org/#!Synapse:syn4894912), included transcriptome data of 376 brain samples from temporal cortex (TCX) and cerebellum of AD patients and control individuals (age-matched healthy individuals and patients with progressive supranuclear palsy, PSP, (41) as an unrelated neurodegenerative disease control). Using an unbiased strategy, we first determined expression values of transposable element (TE) families, restricting the analysis to datasets with high RNA Integrity Number (RIN>7). Global expression analysis of brain samples revealed that upregulated TE families were mostly (∼90%) primate-specific elements (present only in human and primate genomes), while the rest of the detected expression derived from ancient TEs, found also in other mammals/vertebrates (Supplementary Fig. 6a). In the human brain, the most abundantly expressed TEs identified by us were non-LTR elements, such as SINE-VNTR-*Alu*s (SVA) and the retrotranspositionally active L1_Hs (42), followed by endogenous retrovirus groups, HERV-E_a and HERV-K (Supplementary Fig. 6a and b). While investigating TE expression in AD compared to controls, we observed differential HERV-K transcription in TCX but not in cerebellum samples (data not shown). Thus, for detailed analysis, we established a TCX data frame (total *n* = 176) of AD samples (*n* = 82) and a combined control of age-matched healthy and PSP samples (*n* = 94) (Supplementary Fig. 6c). In AD samples, we identified HERV-K as the most expressed and differentially upregulated TE (Fig. 6a and Supplementary Fig. 6c). Of further note, we observed reduced expression of AluY elements in AD samples compared to controls, while, despite their high overall expression, L1 and SVA were not among the differentially expressed TEs (Fig. 6a).

**Fig. 6.**
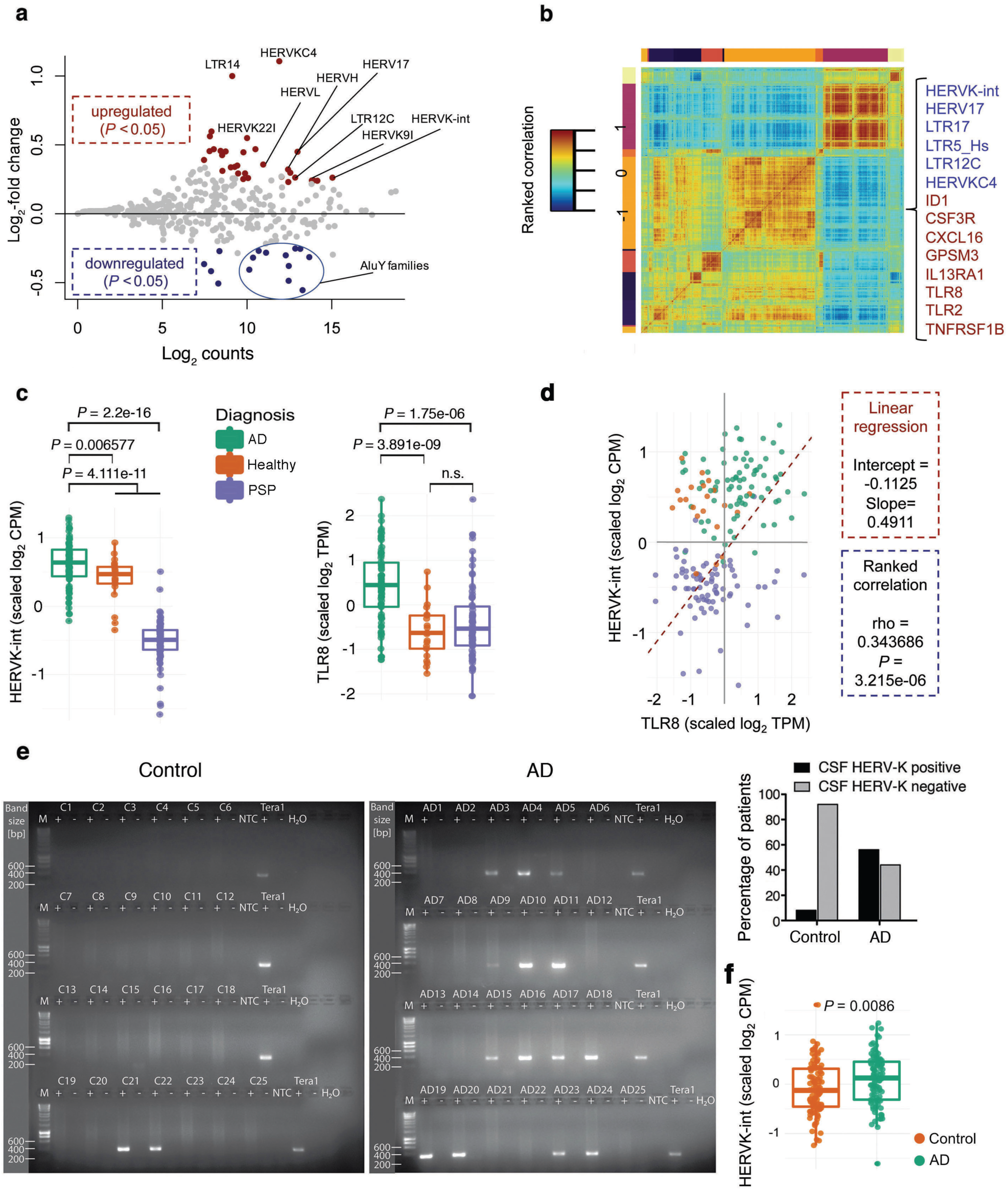
Correlated upregulated HERV-K and TLR8 expression in AD patients. (**a**) Differentially expressed transposable elements (TEs) in the temporal cortex (TCX) of Alzheimer’s disease (AD) patients. MA plot shows differential TE expression (log_2_-fold change) in the TCX of 176 post-mortem samples of AD patients (*n* = 82) *vs*. control individuals (combined ageing healthy individuals and patients with progressive supranuclear palsy (PSP), *n* = 94). Each dot represents one TE family. Dots are coloured if the *P* value from DESeq2/limma statistics was <0.05. Note that the most differentially up- and downregulated TEs in AD patients vs. controls include HERV-K-int (as designated in Repbase; i.e. HERV-K(HML-2); see the paper text), HERVK9I and various AluY subfamilies. (**b**) Correlated expression of HERVK-int (HERV-K(HML-2)) with immune response genes. Clustered pairwise correlation matrix generated by weighted gene coexpression network analysis (WGCNA) across 176 brain samples (see **a**, including ∼2,400 genes and 17 TE families, Spearman rank correlation, Euclidian distance). In a highlighted cluster, TEs (blue) including HERVK-int/LTR5_Hs (HERV-K(HML-2)) are co-expressed with several immune response genes (dark red), including TLR8 and TLR2. (**c**) Jitter boxplots visualize Z-score-transformed, unbiased relative expression of (left) HERV-K-int/HERV-K(HML-2) (log_2_ CPM) and (right) TLR8 (log_2_ TPM) in TCX samples of AD, PSP and healthy ageing controls. Every dot represents one sample. *P* values were determined by Wilcoxon test, which was further adjusted for multiple corrections, as indicated. n.s., not significant. (**d**) Correlation analysis of HERVK-int/HERV-K(HML-2) and TLR8 expression. Scatter plot shows scaled log_2_-transformed expression of HERVK-int/HERV-K(HML-2) (CPM) and TLR8 (TPM) in the analysed samples (colour codes defined as in **c**). Linear regression analysis (dashed line) indicates correlation between HERVK-int/HERV-K(HML-2) and TLR8 expression in AD samples. Rho values were obtained from pairwise ranked correlation analysis. (**e**) Cerebrospinal fluid (CSF) from patients with AD (*n* = 25) and control subjects (*n* = 25) were assayed by endpoint RT-PCR using primers specific for HERV-K *env* region encompassing the TLR7/8 recognition motif (left). Quantitative analysis of the frequency of HERV-K detection in CNS samples from control individuals and AD patients (right). (**f**) Presence of HERV-K(HML-2) in the CSF of AD and age-controlled samples. Jitter boxplots represent the average of Z-score-transformed counts of expressed HERV-K(HML2) loci (*n* = 98) in the CSF of AD (*n* = 6) and control (*n* = 6) samples. Every dot represents a distinct HERV-K(HML-2) locus. *P* value was determined by Wilcoxon test, as indicated.

In order to fetch genes with expression patterns similar to HERV-K, we produced a pairwise correlation matrix generated by weighted gene coexpression network analysis (WGCNA) across the 176 TCX samples described above, including ∼2,400 genes and 17 TE families. WGCNA assigned HERV-K expression to a correlation cluster together with several immune response genes, including TLR8 and TLR2 (Fig. 6b). Both HERV-K and TLR8 expression were upregulated in AD versus control samples (Fig. 6c), and relative to HERV-K TLR8 exhibited the highest correlation value (Fig. 6d), also when compared to statistically still significant values for TLR2, TLR5, and TLR7 (Supplementary Fig. 6d). Reanalysis of further, previously published RNA-seq datasets derived from brains of AD patients and control subjects (GSE53697 (43), GSE104705 (44)) corroborated specific upregulation of both HERV-K and TLR8 expression in AD (Supplementary Fig. 6e).

To investigate whether endogenous HERV-K RNA is released during neurodegenerative processes such as AD, we analyzed cerebrospinal fluid (CSF) of patients with AD and CSF of a control group (for patient details, see Supplementary Table 1) for presence of HERV-K transcripts by RT-PCR. HERV-K RNA was detected in the CSF of 14 out of 25 (56%) AD patients, whereas only 2 out of 25 (8%) healthy control individuals were positive for HERV-K RNA (*P* = 0.0006) (Fig. 6e). Sanger-based amplicon-sequencing of amplified RT-PCR products from CSF samples positive for HERV-K RNA confirmed RT-PCR products as the targeted HERV-K *env* gene region and indicated transcripts with the phylogenetically younger sequence variant 5’-GUUG*C*GU-3’ to be the dominant variant based on peak heights (data not shown). Next, we compared CSF-derived small RNA-seq datasets from AD patients and healthy control individuals (45) with regard to expression of TEs, including HERVs. Small RNA derived from LTR5_Hs/HERV-K was found enriched exclusively in AD patients (Fig. 6f), further supporting an association between HERV-K activation and AD.

## Discussion

The multi-copy human endogenous retrovirus group HERV-K(HML-2) is physiologically present in various loci of the human genome and is transcriptionally active. Several HERV-K(HML-2) (referred to as HERV-K in the following) proviruses in the human genome formed recently and are thus relatively intact, permitting the expression of both viral RNA and proteins (10). Inflammatory and, as yet unidentified, further stimuli in CNS diseases may modulate and trigger endogenous retrovirus re-activation and transcription (39), thus promoting viral RNA expression, which may further modulate the pathological status. Previous studies have suggested that the principal mechanisms by which endogenous retroviruses can injure the brain are based on the release of virus-encoded proteins or the induction of potential immunopathogenic molecules such as TNF-α from infected or activated host immune cells (24, 46). By contrast, our current study indicates a novel functional role for human endogenous retroviral RNA transcripts in neurodegenerative processes. Several molecules have been identified that activate innate immunity when released from injured eucaryotic cells. Although these endogenous activators can be evolutionarily highly conserved molecules, their common feature is being foreign to the extracellular environment in a normal tissue (1). As TLRs recognize exogenous retroviral RNA, we considered human endogenous retroviruses as potential activators of these receptors. On the basis of sequence similarity to known TLR7 and TLR8 ligands derived from exogenous retroviruses such as ssRNA40 from HIV, which triggers neurotoxicity in the brain (47), we asked whether a similar GU-rich sequence present in the HERV-K *env* gene could also activate TLR signaling. Indeed, using both biochemical and biophysical approaches we found that HERV-K-derived RNA containing this GU-rich motif binds directly to human TLR8 and induces signaling through both human TLR8 and murine Tlr7. Based on these findings, we investigated human-derived cell systems in terms of neuronal injury and modelled TLR-mediated neurodegenerative processes by delivering HERV-K RNA extracellularly to neurons from mice that lack HERV-K and instead carry phylogenetically different ERVs (48). Exposure of neurons to HERV-K RNA that harbored GU-rich sequences induced neurodegeneneration through intrinsic murine Tlr7 and human TLR8 signaling. The essential feature of this signal is the mislocalization of an endogenous RNA in the extracellular space, potentially released during cellular injury initiated by a specific disease-causing factor that gains access to TLRs located in neuronal endosomes (5). Accordingly, we found that apoptotic neurons release HERV-K RNA, which in turn induces further neuronal injury. Under physiological conditions, the interaction between these HERV-K RNA transcripts and neuronal Tlr7/TLR8 may be prevented by separated cellular compartments in which these components are located. However, the amount of HERV-K RNA released into the brain parenchyma at the site of injury in neurodegenerative processes and local concentration maxima of such HERV-K RNA are unknown. Concentrations of HERV-K RNA used in the current study were based on our previous work on RNA-mediated neurotoxicity (5, 47), and may be considered supraphysiologic, especially since neurotoxicity observed in our current work occurred within a time period of days and weeks while neurodegenerative processes in the context of CNS disorders may evolve over multiple decades. Thus, our study presents a proof of principle regarding the effects of HERV-K RNA in the brain, and future work will be required to determine pathophysiological concentrations and key properties of HERV-K RNA as a signaling molecule in the brain.

Although we cannot rule out that the observed neurodegenerative effects are not specific for HERV-K RNA, but can be caused by any RNA species containing the GU-rich TLR7/TLR8 recognition motif, and that presence of that motif in HERV-K is incidental, several aspects have to be taken into account when discussing a specific role for HERV-K RNA in this context. First, among HERVs the sequence motifs GUUGUGU and GUUGCGU were detected in HERV-K as a common sequence portion within the *env* gene. Second, several differently sized HERV-K-derived nucleotides (22-nt, 50-nt, and 785-nt derived from the concerned HERV-K *env* region), as well as endogenous, retroviral particle-derived full-length HERV-K transcripts, were capable of inducing neuronal inury. Third, HERV-K RNA is present and stable in CSF. Finally, HERV-K inhibitor reduced the neurotoxic effect of lysates from dying neurons applied to neurons. Given that LNA-based RNA inhibitors, such as the one used in our study, are highly sequence-specific, even at the level of single mismatches (49), our results indicate that HERV-K RNA is indeed involved in the induction and/or progression of neurodegeneration.

As noted, HERV-K harbors the sequence motif 5’-GUUGUGU-3’, which mediates species-specific recognition of Tlr7 and TLR8 in mouse and human, respectively. Substantial reduction of the GU content in this motif abolished Tlr7 activation in neurons in our study. However, the ability to stimulate Tlr7 depended on a GU-rich element rather than on the exact GUUGUGU motif, as HERV-K RNA transcript subsets that contain slight variations in this motif (GUUG*C*GU) effectively engaged Tlr7. HERV-K proviruses harboring GUUG*C*GU are among the evolutionarily youngest proviruses (Supplementary Fig. 1), that is, such proviruses were formed in the genome relatively recently. We found that HERV-K RNA inhibitor reduced the neurotoxic effect of virions known to harbor RNA from several HERV-K loci, including loci containing the TLR7/8 recognition motifs (12, 14). Taking into account the sequence specificity of the LNA-based RNA inhibitor (see above), our data strongly indicate that HERV-K RNA physiologically expressed in human can induce neurodegeneration.

We show that mTlr7 and hTLR8 play a role beyond their function as immune receptors in immune cells, demonstrating that they also serve as death receptors in CNS neurons. Although microglia released inflammatory molecules in response to HERV-K RNA and enhanced HERV-K RNA-induced neuronal cell death, they were not required for the injurious effect of extracellular HERV-K RNA applied to neurons *in vitro*. Since *in utero* electroporation at embryonic stage E14.5 only targets neurons and not glia (50), only those neurons newly expressing mTlr7 or hTLR8 were susceptible to HERV-K RNA, as neighboring neurons lacking these TLRs were unaffected. This result is consistent with our *in vitro* assays suggesting that HERV-K RNA activates an intrinsic cell death pathway in neurons. Further, we demonstrate that neuronal injury induced by HERV-K RNA requires SARM1, a member of the TIR-containing adaptor family. It has been previously identified as a regulator of neurodegeneration in *Drosophila* and mice, indicating SARM1 as a member of an ancient neuronal death signaling pathway (33). Virus-induced RIG-1/MAVS signaling as well as TLR7/TLR9 signaling cause mitochondrial accumulation of SARM in neurons (35, 51), which triggers intrinsic apoptosis by generating reactive oxygen species and depolarizing the mitochondrial potential (52). Furthermore, SARM1 acts as a negative regulator of TLR-mediated canonical signaling pathways, including NF-κB activation (53, 54). Accordingly, we found that HERV-K RNA does not induce NF-κB activation in neurons, and Myd88 is not involved in HERV-K RNA-mediated neurodegeneration. However, the exact signaling cascade linking Tlr7 and TLR8 signaling with caspase-3 activation in HERV-K-triggered neurodegeneration remains to be elucidated.

HERV expression has been associated with several neurological diseases, although an etiologic demonstration that HERVs cause or contribute to the particular diseases is lacking so far. Our analysis of transcriptome datasets derived from a large cohort of AD patients and control individuals identified substantial correlation of upregulated HERV-K and TLR8 expression (TLR7 expression being less correlated) in brains of AD patients compared to control subjects. Notably, this correlation was specifically observed in the temporal cortex, but not in cerebellum. Also, we observed that the most active TEs in the human temporal cortex are phylogenetically young, primate-, and/or human-specific HERV-derived elements (e.g. HERV-E, LTR5_HS/HERV-K, SVA) and some potentially mutagenic LINEs from the L1_Hs, L1_PA2, and L1_PA3 subfamilies. SVA and HERV-K can form active chromatin regions in the brain and thus affect neighbor gene expression. As they are polymorphic in the human population, they might be considered as factors in susceptibility to human-specific neurological diseases, such as AD (55, 56). In agreement with previous reports (57), we detected an overall high expression of potentially mutagenic L1_Hs elements in the tested brain samples. However, unlike HERV-K, L1_Hs was not differentially upregulated in AD patients. Also, HERV-K RNA encompassing the TLR7/8 recognition motif was detected more often in CSF of AD patients (56% of samples) than in control individuals (8%) by PCR, and these data were corroborated by reanalysis of small RNA-seq datasets derived from AD patients. Thus, our data may have implications for AD patients with increased HERV-K levels in CSF. Moreover, since HERV-K RNA can induce CNS injury, blocking functional effects of HERV-K transcripts may affect the course of AD. In our study, a HERV-K RNA-specific inhibitor protected neurons from neurotoxic effects induced by HERV-K transcripts. The HERV-K inhibitor also prevented neurodegenerative effects and increasing microglial numbers in response to HERV-K in *APPPS1* mice. Nevertheless, our observation of neurodegeneration and microglial accumulation, but no impact on the Aβ plaque burden, in response to intrathecal HERV-K RNA in the *APPPS1* AD mouse model points to a more complex role for HERV-K in AD, suggesting that transgenic mouse models such as the *APPPS1* only partially recapitulate specific disease aspects such as cerebral amyloidosis and basic disease mechanisms (58).

To the best of our knowledge, we present here the first conclusive evidence for an injurious role of HERV-K-derived transcripts in the brain. We speculate that RNA derived from ERVs is released under as yet undetermined pathological conditions and stimulates TLRs, further accelerating neuronal decay, as it may occur in neurodegenerative diseases. Consistent with this hypothesis, HERV-K transcripts were found more frequently in the CSF of AD patients as in CSF from control individuals. Thus, it appears that stable extracellular HERV-K RNA in human CSF exists that is capable of stimulating membrane receptors. However, further research including more differentiated and larger patient cohorts will be required to investigate the probably more complex role of HERV-K in the CNS, to establish clinical consequences of neurodegeneration triggered by HERV-K through TLRs in specific CNS disorders such as AD, and to identify means to prevent neurodegenerative consequences from HERV-K.

## Methods

### Reagents

The following RNA oligoribonucleotides, in which “s” depicts a phosphothiorate linkage, were synthesized by Purimex DNA/RNA-Oligonucleotide, Grebenstein, Germany, or Integrated DNA Technologies, Leuven, Belgium: 22-nucleotide HERV-K(HML-2) (referred to as HERV-K or HERV-K (+GU) in the following), 5’-CsCsUsUsUsAsCsAsAsAsGsUsUsGsUsGsUsAsAsAsGsC-3’, HERV-K upstream of GUUGUGU motif, 5’-AsAsUsCsUsCsUsAsCsCsCsCsAsAsGsAsCsCsAsAsAsA-3’ (referred to as HERV-K (-GU)), 50-nucleotide HERV-K, 5’-GsUsCsUsAsAsCsAsGsUsUsCsCsUsUsUsAsCsAsAsAsGsUsUsGsUsGsUsAsAsAsGsCsCsCs CsCsUsUsAsUsAsUsGsCsUsAsGsUsU-3’ (referred to as HERV-K long), scrambled 50-nucleotide HERV-K, 5’-UsUsGsGsGsAsUsUsCsCsUsAsAsUsAsAsUsUsCsAsGsUsAsAsUsCsUsAsCsUsCsCsUsUsUs CsCsAsAsCsUsGsAsUsGsGsGsCsAsU-3’ (referred to as HERV-K long mutant), oligoribonucleotides with modified sequence, 5’-GsGsUsCsAsUsAsCsUsGsUsAsGsAsAsUsUsAsAsCsCsU-3’ and 5’-UsGsAsGsGsUsAsGsUsAsGsGsAsAsGsUsGsUsGsGsUsA-3’ (referred to as control oligoribonucleotide #1 and #2, respectively), mutant oligoribonucleotide, 5’-UsGsAsGsGsUsAsGsAsAsGsGsAsUsAsUsAsAsGsGsAsU-3’, HERV-K young, 5’-CsCsUsUsUsAsCsAsAsAsGsUsUsGsCsGsUsAsAsAsGsC-3’. For some experiments, RNA oligoribonucleotides without phosphorothioate linkages were used. Loxoribine, R848, TL8-506, and polyinosine-polycytidylic acid were purchased from InvivoGen (San Diego, USA). LPS was obtained from List Biological Labs (Campbell, USA).

### Mice and cell lines

C57BL/6 (wild-type) mice were obtained from the FEM, Charité – Universitätsmedizin Berlin. Tlr7KO, Myd88KO, and Tlr2/4KO mice were generously provided by S. Akira (Osaka University, Japan). Sarm1KO and *APPPS1* mice were maintained, as previously described (38, 59). Immortalized bone marrow-derived macrophages were generated as previously described (60). Tera-1 cells, derived from a lung metastasis of a human embryonal testicular carcinoma, were maintained, as previously described(12), in McCoy’s 5a medium containing 10% heat-inactivated FCS and penicillin (100 U/ml)/streptomycin (100 µg/ml). SH-SY5Y neuroblastoma cells (provided by Dr. Markus Höltje, Charité –Universitätsmedizin Berlin), HEK 293 cells and HEK-Blue^TM^ cells transfected with murine or human TLR7 or murine or human TLR8 as well as the respective control cell lines HEK-Blue^TM^ Null1, Null1-v, Null1-k and Null2-k (all obtained from InvivoGen) were cultured in DMEM supplemented with 10% heat-inactivated fetal calf serum and penicillin (100 U/ml)/streptomycin (100 μg/ml). THP-1 cells (provided by Dr. Elisabeth Kowenz-Leutz, Max Delbrück Center for Molecular Medicine, Berlin) were cultured in RPMI-1640 medium, supplemented with 0.05 mM 2-mercaptoethanol, 10% heat-inactivated fetal calf serum and penicillin (100 U/ml)/streptomycin (100 µg/ml). Four days prior to usage, THP-1 cells were differentiated into macrophages by incubation with 100 ng/ml phorbol 12-myristate 13-acetate (PMA) for 48 h. After differentiation, a complete media change was performed, and cells were cultured for another 2 d in PMA-free media. Cells were grown at 37°C in humidified air with 5% CO_2_.

### Immunocytochemistry and immunohistochemistry

Immunostaining was performed as described previously (4). We used primary antibodies against NeuN, Neurofilament, Microtubule-Associated Protein 2 (all 1:1,000, Millipore, Burlington, USA) and active caspase-3 (BD Pharmingen and Cell Signaling Technology, Danvers, USA, 1:300). Iba1 antibody was purchased from Wako Chemicals, Neuss, Germany (1:500). IB4 was obtained from Invitrogen, Carlsbad, USA, and DAPI was purchased from Roche, Basel, Switzerland. (Fluorescence) microscopy was done using an Olympus BX51 microscope and a Leica TCS SL confocal laser scanning microscope with sequential analysis (argon, 488 nm; helium neon, 543 nm).

Aβ immunostaining was carried out with the primary antibody 4G8 (Bio Legend, San Diego, USA). Sections were pre-treated with 0.3% H_2_O_2_ in PBS (15 min), washed in H_2_O_dd_ (5 min), then heated to 95°C in 0.01 M citrate buffer pH 6.0 (∼2 min + 15 min cool-down), treated with PBS/0.1% Triton X-100 (15 min) and incubated in 88% Formic Acid (3 min). After 2x washing in PBS (5 min), sections were blocked for 1 h in PBS supplemented with 10% fetal bovine serum (FBS)/4% fat-free milk powder. Primary antibody (dilution 1:1000) was applied in PBS/10% FBS (16 h) and washed in PBS/Triton X-100 (5 min) and PBS (1 min). Sections were incubated with biotinylated secondary antibody (Agilent, diluted 1:500 in PBS/10% FBS) for 1.5 h, followed by washing in PBS/Triton X-100 and PBS for 1 min. DAB was developed for 90 s using the Vectastain Elite ABC Kit (Vector Laboratories, Burlingame, USA) according to the manufactureŕs instructions.

### Toxicity assays *in vitro*

For toxicity studies, the indicated amounts of RNA oligoribonucleotides and other reagents were directly added to neuronal cultures for time periods as indicated in the respective figures. Control cultures were incubated with unmodified media. LPS (100 ng/ml for microglia, 10 μg/ml for neurons) served as a positive control for TLR4 activation and microglia-mediated neurodegeneration in microglia/neuron co-cultures. Loxoribine (1 mM) served as a positive control for TLR7-mediated effects, while TL8-506 (10 μg/ml) served as a positive control for TLR8 activation. Inhibition of HERV-K transcripts was achieved by using an inhibitor oligoribonucleotide with full phosphorothioate backbone (LNA, custom-designed and provided by Exiqon, Vedbaek, Denmark). The strand was: 5’-TACACAACTTTGTAAA-3’. A nonspecific inhibitor oligoribonucleotide (5’-ACGTCTATACGCCCA-3’) was used as negative control inhibitor (Exiqon). Neurons were seeded in 24-well-plates (5 × 10^5^ per well), and after 24 h, cells were incubated with the HERV-K inhibitor (10 nM) or the nonspecific inhibitor in the presence of virion particles isolated from Tera-1 cells. Cells were immunostained with NeuN antibody after 6 d. For each condition, experiments were performed in duplicates. NeuN-positive cells were counted by analyzing six high-power fields per coverslip. The viability of control cells was set to 100%. The numbers of NeuN-positive cells observed for each condition were compared with control conditions, and results were expressed as relative neuronal viability.

### Intrathecal injection into mice

Intrathecal injection of C57BL/6J mice was performed as described previously (61). We used 10 µg RNA for injections. For HERV-K inhibition experiments, 125 pmol of HERV-K inhibitor (see above) or the nonspecific inhibitor were injected intrathecally 16 h before the intrathecal injection of the respective HERV-K or the mutant oligoribonucleotide. Brains were analyzed after 3 d or 2 weeks, as indicated in the respective figure.

At 4 weeks of age, *APPPS1* mice were intrathecally injected with 10 µg HERV-K RNA combined with or without 125 pmol HERV-K inhibitor treatment given intrathecally 16 h beforehand. Sham-operated mice received PBS instead of the oligoribonucleotide intrathecally. Subsequently, injections were repeated every 4 weeks until the end of the experiment (120 d of age).

After transcardial perfusion with 4% paraformaldehyde, brains were surgically removed and cryoprotected in 30% sucrose. Neuronal survival and numbers of microglia were analyzed by blinded quantification counting cortical NeuN-/active caspase-3-positive cells and Iba1-positive cells, respectively, in six fields (magnification x60) of five representative sections of each brain. For plaque load quantification (4G8), we sampled 3 series of 5 coronal sections per animal with a total of ∼60 non-overlapping images of the cortical plaque burden at 200 x magnification (fixed exposure/illumination). Histogram data for individual mice were retrieved from these images using ImageJ 1.48. Statistical plaque load analysis was carried out based on obtained histogram data, and expressed as average percentage of stained pixels at a fixed threshold for each animal.

### Plasmids and *in utero* electroporation

The plasmid pCAG-mTLR7-IRES-EGFP was constructed by subcloning the mTlr7 coding sequence from pUNO-TLR7-HA (Invitrogen, Carlsbad, USA) into pCAG-IRES-EGFP, as previously described (5). The plasmid pCAG-hTLR8-IRES-EGFP was generated by blunt ligation of pUNO-hTLR8 (InvivoGen, San Diego, USA) digested by *Age*I and *Nhe*I into *EcoRV* digested pCAG-IRES-EGFP. pCAG-IRES-tdTomato served as control vector. Plasmids were purified using an endotoxin-free plasmid purification kit (Qiagen, Hilden, Germany) and eluted in nuclease-free water. Prior to electroporation, plasmids were diluted to 2 µg/µl with water and mixed with 1 µl Fast-Green per 20 µl DNA. *In utero* electroporation was carried out, as described (62) with minor adjustments. Briefly, timed pregnant mice at E14.5 (post coitum) were deeply anesthetized and kept on a warming pad (31-32°C) during the entire surgical procedure. The anesthesia was maintained by a steady supply of isoflurane (DeltaSelect) and oxygen (1 L/min). The uterus with the embryos was exposed performing a ∼2 cm midline incision in the ventral abdomen peritoneum and moistened with warmed PBS containing antibiotics (1000 U/ml penicillin, 1000 µg/ml streptomycin). Plasmid DNA was injected into the lateral ventricle using a glass capillary and with the aid of a pico spritzer. Electroporation was carried out by applying six electric square pulses of 35 V each, 50 ms duration and 950 ms intervals using an ECM830 electroporator (BTX Harvard Apparatus). Subsequently, the uterus was placed back into the abdominal cavity, and the body wall was sutured. One third of the litter received the control (tdTomato) plasmid. P19 animals were used for intrathecal injection of RNA, as described above.

### Overexpression of endogenous HERV-K by lentiviral transduction

We utilized lentiviral overexpression in SH-SY5Y cells for transcription of HERV-K RNA. Within a lentiviral shuttle vector (FUGW) (63), a 785-nt HERV-K sequence portion (corresponding to nt 7291-8076 in Genbank accession number AF074086.2 (64)) was cloned downstream of a nucleus-targeting GFP (NLS-GFP) reporter gene that was controlled by a strong synthetic CAG promoter. The HERV-K sequence contained either the TLR7 recognition motif 5’-GTTGTGT-3’ f(CAG)-NLS-GFP-HML2(TLR)-w or an engineered mutated motif 5’-GTTG*C*GT-3’ (f(CAG)-NLS-GFP-HML2(noTLR)-w). As a further control, HERV-K sequences with wild-type or mutated TLR7 recognition motif were cloned in reverse-complement orientation into the lentiviral vector (f(CAG)-NLS-GFP-asHML2(TLR)-w, f(CAG)-NLS-GFP-asHML2(noTLR)-w). Viral production was performed by the Charité Viral Core Facility (http://vcf.charite.de), as described previously (63). The following lentivirus constructs were used: BL-1177 f(CAG)-NLS-GFP-HML2(TLR)-w; BL-1178 f(CAG)-NLS GFP-HML2(noTLR)-w; BL-1179 f(CAG)-NLS-GFP-asHML2(TLR)-w; BL-1180 f(CAG)-NLS-GFP-asHML2(noTLR)-w.

### Primary culture of cortical neurons and microglia

Primary cultures of purified cortical neurons and microglia, as well as the respective co-cultures were generated as described previously (3).

### Isolation of virions from Tera-1 cells

Isolation of viral particles released from Tera-1 cells (12, 14, 65) into the culture media was performed using Total Exosome Isolation Kit (Invitrogen) according to the manufacturer’s instruction. Briefly, cell culture supernatant was centrifuged at 2,000 g for 30 min at 4°C. 0.5 volume of exosome isolation reagent was added to the cell-free culture media and mixed by vortexing before incubating overnight at 4°C. Samples were centrifuged at 10,000 g for 1 h at 4°C. Supernatant was removed. The pellet containing virions was re-suspended in PBS and was used for toxicity assays *in vitro*.

### RNA extraction from cell culture, supernatant and CSF

Total RNA from cells was isolated using the RNeasy Mini Kit (Qiagen) following the manufacturer’s protocol and using 5 x 10^6^ cells in total. After centrifugation of cells at 300 x g for 5 min the supernatant was completely removed and 600 µl disrupting buffer was added to the cell pellet. The lysate was homogenized using the QIAshredder spin columns (Qiagen). 1 volume of 70% (v/v) ethanol was added to the homogenized lysate and then transferred into an RNeasy spin column. After three washing steps, RNA was eluted by two centrifugation steps using 40 µl RNase-free water.

Isolation of total RNA from supernatant of cultured cells was performed using the *mir*Vana^TM^ Paris^TM^ RNA Purification Kit (Thermo Fisher Scientific, Waltham, USA), following the manufacturer’s protocol. Supernatant was centrifuged (10 min x 1,200 g) prior to the isolation procedure to remove all cell remnants. Subsequently, an equal volume of 2x denaturating solution was added to the cell-free supernatant and incubated on ice for 5 min. One volume of acid-phenol:chloroform was added to the mixture, vortexed for 60 s and centrifuged for 5 min. The aqueous phase was transferred and mixed with 1.25 volumes of pure ethanol and mixed thoroughly. Step-by-step, the whole mix was pipeted onto a filter cartridge, which was then washed 3x. RNA was eluated with 95°C water.

For RT-PCR analysis of CSF, two groups of subjects were selected from the Kompetenznetz Demenzen biomaterial bank. The control group comprised healthy subjects with subjective memory complaints. The second group consisted of patients diagnosed with Alzheimer’s disease. Neuropsychometric and biochemical characteristics as determined in the CSF were consistent with the diagnosis of either group. For total RNA isolation from CSF the QIAamp Viral RNA Mini Kit (Qiagen) was used according to the manufacturer’s protocol. Briefly, 500 µl CSF were mixed with one equal volume of lysis buffer and incubated at RT for 10 min, then loaded onto the QIAamp Mini column. After two washing steps, RNA was eluted using 40 µl of AVE Buffer at each step.

Following RNA isolation, all RNA preparations were DNase-treated using Turbo DNA-free Kit (Invitrogen) to remove contaminating DNA according to the manufacturer’s protocol. 10 µg of total RNA were incubated with 1 μl Turbo DNase in 1x reaction buffer for 30 min at 37°C. Subsequently, 0.1 volume of thoroughly resuspended DNase Inactivation Reagent was added to the mixture and incubated for 5 min at room temperature followed by centrifugation at 10,000 x g for 1.5 min. Supernatant was transferred to a new tube and used for PCR analysis and toxicity assays.

### HERV-K-specific RT-PCR

11.5 µl of RNA isolated from CSF was used for reverse transcription PCR in a 20 µl reaction volume using the Moloney Murine Leukemia Virus Reverse Transcriptase (M-MLV RT; Promega, Madison, USA). The RT mixture was incubated for 60 min at 37°C followed by an inactivation step for 10 min at 70°C. With each sample, an RT+ reaction (including MMLV-RT) and an RT-reaction (without MMLV-RT) were set up from the same mastermix. Presence of HERV-K RNA was evaluated by endpoint RT-PCR using the GoTaq DNA Polymerase (Promega, Fitchburg, USA) and 1 µl of the generated cDNA in a 20 µl total volume reaction following the manufacturer’s protocol. Each reaction mixture contained 0.5 µM of HERV-K *env*-specific primers (HERV 101F: 5’-TCTACCCTTGGGAATGGGGA-3’; HERV 499R: 5’-AGCAGAATACGGTGTTGCCA-3’). Amplification was performed using a BioRad C1000 Touch Thermal Cycler at 95°C for 5 min followed by 40 cycles of 95°C for 50 s, 48°C for 50 s, 72°C for 1 min and a final elongation step for 10 min at 72°C. 10 µl of the amplification product was run on a 1.5% agarose gel.

### Multiplex immunoassay

Microglia, bone marrow-derived macrophages, or macrophages differentiated from THP-1 cells were incubated for various durations, as indicated in the respective figures, with increasing concentrations of HERV-K or control oligoribonucleotides complexed to the transfection agent LyoVec (InvivoGen). Subsequently, supernatants were evaluated using the ProcartaPlex Multiplex Immunoassay (Invitrogen) following the manufacturer’s protocol. Briefly, 50 µl of magnetic capture beads were plated and washed before adding 50 µl cell culture supernatant, centrifuged at 4°C for 10 min at 1,000 x g, and incubated with 50 µl of assay buffer overnight. Subsequently, detection antibody mixture was added to the wells before washing and adding the reading buffer to prepare the samples for running on a Bio-Plex System (Bio-Rad, Hercules, USA).

### Flow cytometry

After fixation and permeabilization (Cytofix/Cytoperm Kit BD Biosciences, Heidelberg, Germany), SH-SY5Y cells and primary murine neurons were incubated for 30 min at 4°C, using the following antibodies: fluorescein isothiocyanate (FITC)-conjugated anti-TLR7 (Imgenex, San Diego, USA), Phycoerythrin (PE)-conjugated anti-TLR8 (Abcam, Cambridge, UK), Alexa Fluor® 647 anti-β-Tubulin (eBioscience, San Diego, USA), and their recommended isotype controls (Imgenex or eBioscience, San Diego, USA) (all antibodies were used at 1:100). Blocking of Fcγ-receptors (eBioscience) was performed before cell surface and intracellular staining. Flow cytometric analysis was performed on FACS Canto II (BD Biosciences) and analyzed by FlowJo software (TreeStar, Inc.). TLR expression results were quantified by median fluorescence intensity and presented in histograms.

### Electrophoretic mobility shift assay

Electrophoretic mobility shift assays were performed as described previously (66).

### HEK-Blue reporter assay

One day after seeding 50,000 cells/well in a 96-well-plate, HEK-Blue SEAP reporter 293 cells expressing mTlr7, mTlr8, hTLR7 or hTLR8 and the inherent control cells were transfected with different concentrations of HERV-K or control oligoribonucleotides, as described above. Cells were stimulated with indicated agents dissolved in 90% HEK-Blue Detection reagent and 10% culture media. Detection of the reporter protein SEAP was performed at 655 nm using Varioskan Flash (Thermo Fisher Scientific). For each condition, 4 wells were individually measured, and the average was used for evaluation.

### Microscale thermophoresis (MST)

Purified polyhistidine-tagged TLR8 protein was purchased from LSBio (Seattle, USA). The protein was delivered in Tris/HCL buffer with 50% glycerol. Receptor labeling was performed following the manufacturer’s protocol (RED-tris-NTA dye, NanoTemper Technologies GmbH, Munich, Germany). TLR8 (2µM) was diluted 1:10 in MST buffer (50 mM Tris/HCl pH 7.4, 150 mM NaCl, 10 mM MgCl_2_, 0.05% polysorbate (Tween) 20, 0.06% n-Dodecyl-beta-Maltoside) to a final concentration of 200 nM. RED-tris-NTA was dissolved in 1x PBS-T buffer to a concentration of 100 nM, mixed 1:1 with TLR8 and incubated on ice for 30 min. MST measurements were performed in the cold-room under controlled ambient temperature of 10°C. To prevent nucleotide oligomerization each oligoribonucleotide was incubated for 5 min at 80°C and placed on ice until the experiment. RNA titration was created with a serial dilution of RNA with water. For each single measurement 5 µl of RNA were mixed with 5 µl RED-tris-NTA-labeled TLR8 and incubated on ice for 5 min shortly before the sample was loaded to standard glass capillaries. Instrumentation and experimental settings: Monolith NT.115 MST_power_ = medium; LED power = 100%. The RED-tris-NTA labeling kit and capillaries were purchased from NanoTemper Technologies GmbH. Data analysis was performed with NanoTemper MO Affinity Analysis V2.3 software and visualized using SigmaPlot V13.0.

### RNA-seq data analysis

Multiple RNA-seq datasets were analyzed for gene and transposable element (TE) expression. The *Mayo* dataset at https://www.synapse.org/#!Synapse:syn4894912 contains data from 376 individuals. Datasets consisting of cerebellum and temporal cortex (TCX) samples derived from Alzheimer’s disease (AD) patients, progressive supranuclear palsy (PSP) patients and elderly healthy individuals, as well as covariate files containing clinical information, were downloaded from the Synapse portal. Subsequently, data were converted from “*bam*” to “f*astq*” format using *bamtofastq* built-in function from bedtools. Following quality control, 45 samples were excluded from the analysis due to low quality (RNA integrity number (RIN) equal or <7). In addition, RNA-seq datasets from different sources were analyzed: dorsolateral prefrontal cortex samples from 9 AD patients and 8 healthy control individuals (GSE53697) (43), 8 samples from the lateral temporal lobe of young healthy individuals (Young Healthy), 10 samples from the lateral temporal lobe of healthy aged individuals (Old Healthy), as well as 12 samples from the lateral temporal lobe from aged AD patients (GSE104705) (44). These data were downloaded as raw sequence reads in *sequence read archive* format and converted to *fastq* files employing SRA toolkit. Due to their variable quality scores, 2-5 terminal nucleotides from the sequencing reads depending on qc scores of *fastq* files were removed. To analyze Refseq genes and TEs simultaneously, coordinates of TE loci were downloaded from UCSC Genome Browser. For further analysis only those TE loci were considered that are not in the vicinity of coding sequences (|5 KB|). Gene track format (gtf) of protein-coding genes and TEs were combined into one gtf file. To estimate gene expression levels, we first indexed the soft-masked hg19/GRCh37 genome providing the transcriptome model (combined gtf of Refseq genes and TEs) using STAR default parameters. For read mapping, we used our defined settings (i.e. *–alignIntronMin 20 – alignIntronMax 1000000 –chimSegmentMin 15 –chimJunctionOverhangMin 15 – outFilterMultimapNmax 20*) for STAR splice mapper (67). Reads with low MAP quality score (<30) were not used. To obtain uniquely mapped read counts of genes and TEs with RefSeq annotations we used *featureCounts*. For TE expression, multimapping reads only counted if they mapped exclusively within a TE family. Gene expression levels were calculated as transcript per million (TPM) counts over a gene (defined as any transcript located between transcription start and end sites). To analyze differential expression of TEs, we counted uniquely mapped reads over given co-ordinates and calculated counts per million (CPM) of mappable reads. Reads per kilobase per million (RPKM) values were calculated as average expression of values for both TE loci and genes only if a pairwise comparison was performed. To calculate differential gene expression, we used the GFOLD algorithm that calculates the normalization constant and variance to extract fold-changes from unreplicated RNA-seq data. To calculate log_2_-fold changes and empirical Bayes-corrected *P* values for each expressed TE family or locus, we subjected normalized counts (using DESeq2) to *limma/Voom* analysis. Note that for comparative analyses datasets from different layouts were never merged into one dataframe. Differential expression of HERV-K was statistically significant between AD and control group analyzing TCX samples, but not cerebellum samples. After this filter step, the TCX dataframe consisted of 82 AD and 94 control (combined PSP and age-matched healthy individuals) samples (40). Of note, our analysis reproduced results previously published by Frost and colleagues (57). Also, TE expression was analyzed with regard to evolutionary age of TE families. To this end, the curated TE classification from *Repbase* was modified as follows: TEs of New World monkeys (Platyrrhini), Old World monkeys (Catarrhini) and hominoids (Hominoidae) were merged into one group as “primate-specific”. Because not annotated entirely correct in *Repbase*, HERV-H was moved from the “eutherian” to the “primate-specific” group. To compare relative expression of an individual TE family or a gene across the dataframe in a pairwise manner, the row-wise Z-score of DESeq2-normalized counts was used, and significance was calculated using the Wilcoxon test.

Small RNA-seq data generated from cerebrospinal fluid (CSF) samples of AD patients and control subjects were provided by Julie A. Saugstad (Oregon Health & Science University, Portland, Oregon) in *fastq* format. Detailed data information is available on the exRNA Research Portal (http://exRNA.org/) and ref. (45). We analysed small RNA-seq raw data by mapping high quality reads (MAPQ >30) to the human reference genome (as described above) using *bowtie2* with the following parameters: --local -p 8 -q --phred33 -D 20 -R 3 -N 0 -L 8 -i S,1,0.50. hg19 co-ordinates annotated by Repeatmasker showing >5 uniquely mapped reads were reported.

### Pathway analysis of differentially expressed genes

Canonical pathways and biological functions of identified differentially expressed genes in datasets were investigated using KEGG pathway and GO tools. Overrepresentation of a biological pathway was assessed by Fisher’s exact test and corrected for multiple testing using the Benjamini-Hochberg procedure. The ratio (overlap) was calculated as the number of genes from the dataset that map to a particular pathway divided by the number of total genes included in the pathway.

### Gene correlational network analysis

To identify significant correlation gene networks, we applied weighted gene co-expression network analysis (WGCNA). This algorithm was employed to determine gene sets co-expressed in the transcriptome of the 176 TCX samples (see above) and identified ∼2400 genes and 17 TE families, including HERV-K, as highly expressed. The metamodel identified probable significant networks. In a significant co-expression module, HERV-K was detected and was then used as a probe to identify HERV-K-associated co-expression clusters. Genes that were correlated beyond the provided correlation thresholds were extracted from the HERV-K cluster. Finally, a pairwise-ranked correlation matrix of the identified genes and TEs was generated. Row-wise Z-scores of DESeq2-normalized counts were displayed as scatterplots, significance of differential expression was calculated using the Wilcoxon test, and rho value and significance of correlation was calculated using *cor.test* built-in R function.

### Statistics

Data are expressed as mean ± s.e.m. or ± s.d., as indicated. Statistical differences over all groups were determined using the nonparametric Kruskal-Wallis test or the one-way ANOVA test, as indicated. Statistical differences between selected groups were determined by the Kruskal-Wallis test with Dunn’s post hoc analysis or the one-way ANOVA test with Bonferroni’s post hoc analysis. Statistical differences between patients with Alzheimer’s disease and control individuals (Fig. 6, Supplementary Fig. 6) were determined by using Student’s *t* test, Fisher’s exact test, or Wilcoxon test, as indicated in figure legends. Statistical differences were considered to be significant at *P* <0.05 for tests with one comparison. According to the Bonferroni’s correction for multiple testing, significance was assumed when *P* <0.05 / (number of comparisons).

### Study approval

All animals were maintained according to the guidelines of the committee for animal care. All animal procedures were approved by the Landesamt für Gesundheit und Soziales (LaGeSo) Berlin. Studies on patients and control individuals have complied with all relevant ethical regulations and were approved by institutional review boards (Ethikkommission Ethikausschuss 1 am Campus Charité-Mitte, Dementia competence network, EA1/182/10; BIH CRG 2a, EA2/118/15). Written informed consent was obtained from all participants prior to inclusion in the study.

## Author contributions

S.L., J.M., Z.I., and K.R. conceived the study and wrote the manuscript. M.C., O.P, D.G., G.K., P.S., V.T., F.H., T.W., and C.S. planned the experiments. P.D., A.N., M.S., M.H., M.S., C.K., K.D., C.S., R.A., O.D., B.C.R., and T.T. planned and carried out the experiments.

## Acknowledgments

This work was supported by Deutsche Forschungsgemeinschaft (DFG) LE 2420/2-1, SFB-TRR167/B3, NeuroCure Exc 257 (to S.L.), SFB740/B6, SFB1078/B6 (to P.S.), DFG Cluster of Excellence ‘Unifying Concepts in Catalysis’ (EXC 314 - Research Field D/E) (to G.K. and P.S.), and European Research Council, ERC Advanced (ERC-2011-ADG 294742) (to Z.I.). J.M. is supported by DFG.

Published data in raw format were provided by Kendall Van Keuren-Jensen, Translational Genomics Research Institute, Phoenix, USA, and Julie A. Saugstad, Oregon Health & Science University, Portland, Oregon, USA. All data associated with this study are available in the main text or Supplemental data.

