## Supplemental for "Human endogenous retrovirus HERV-K(HML-2) RNA causes neurodegeneration through Toll-like receptors"

**Supplemental data**

**Supplementary Figure 1, see next page for legend**

**a**

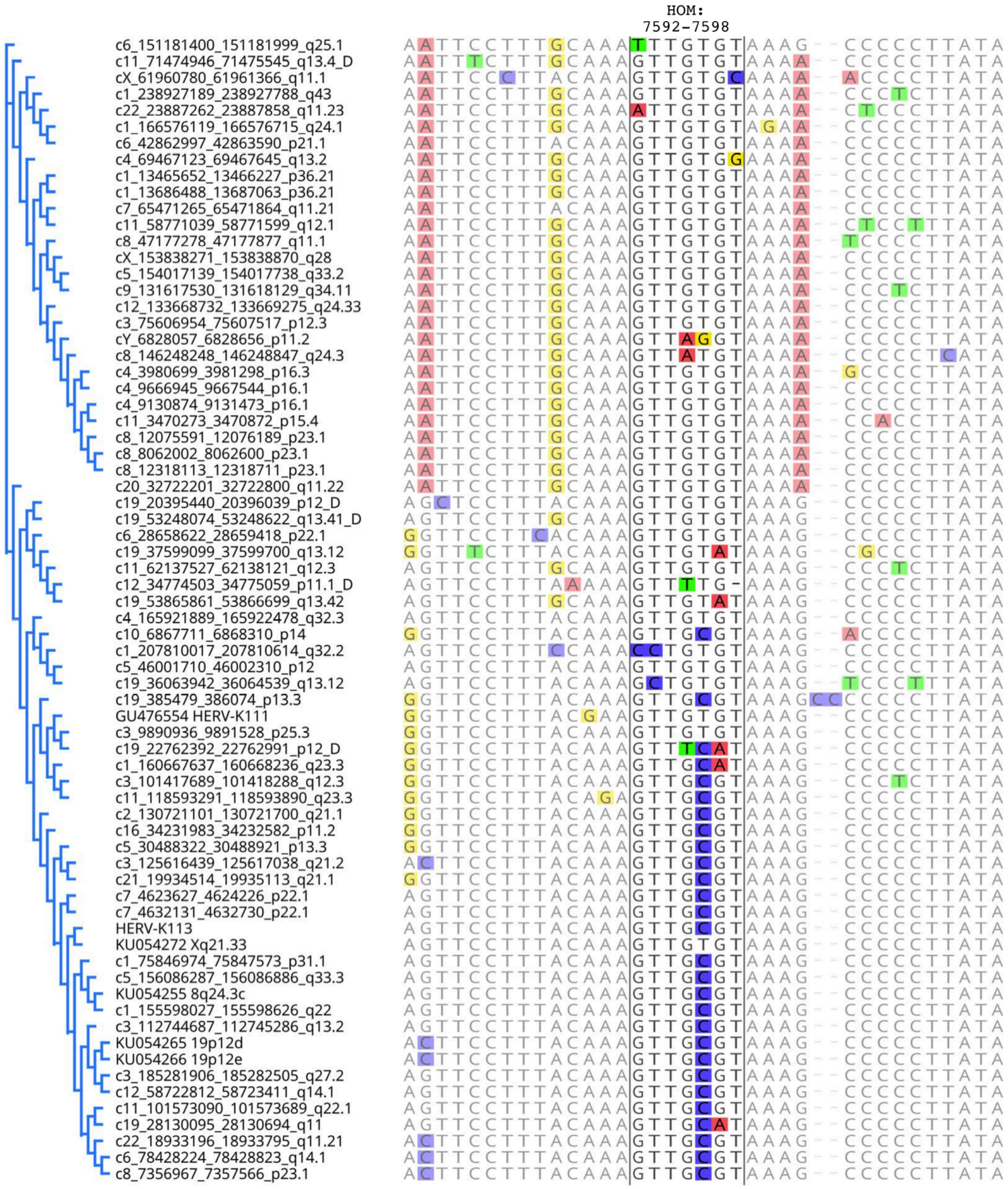

**b**

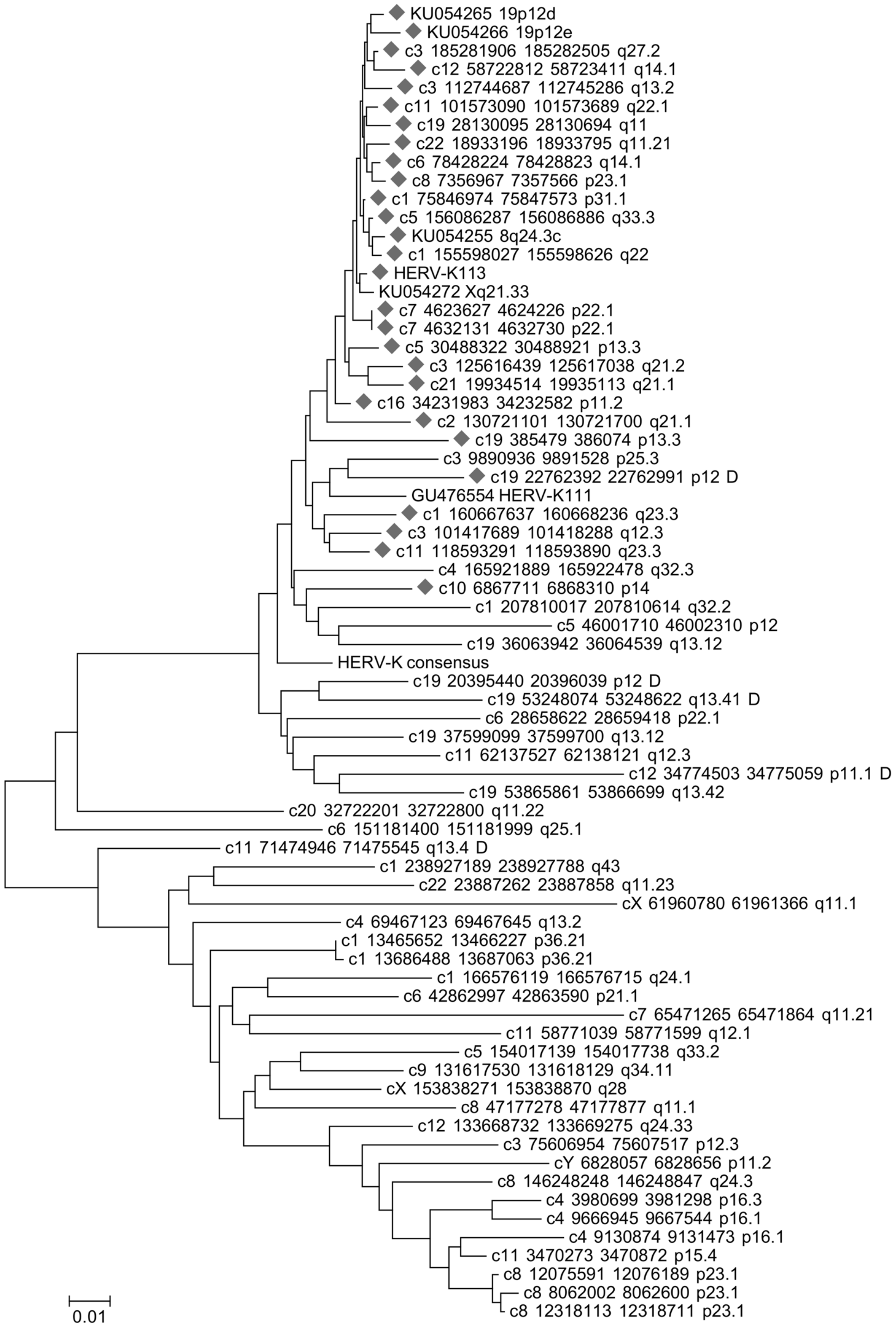


**Supplementary Figure 1. Neighbor-joining tree of HERV-K(HML-2) sequences.** A 200 bp sequence encompassing the TLR recognition motif (GTTGTGT) within the *env* gene region (see the main paper text) was extracted from HERV-K(HML-2.HOM) provirus sequence (Genbank acc. no. AF074086) and used as probe for a BLAT search of the human hg19 reference genome at UCSC Genome Browser. Sequences corresponding to identified chromosome coordinates were extracted with an additional 200 bp of reference sequence added to each end. Sequences were multiply aligned using MAFFT. Only sequences harboring the TLR recognition motif region were included in the alignment. A subregion of the multiple alignment depicting the TLR recognition motif is shown in (**a**). The TLR recognition motif is highlighted and corresponds to nt 7592-7598 in the HERV-K(HML-2.HOM) sequence. Note the variation GTTG*T*GT/GTTG*C*GT (see the main paper text). The cladogram on the left depicts sequence similarities when regarding the entire multiple alignment. (**b**) A neighbor-joining tree was constructed using MEGA7 and the Kimura-2-parameter method. Some of the retrieved sequences harbored internal deletions or insertions of other repeat sequences within the regarded *env* gene region. Such unalignable sequence portions were deleted prior to tree generation (sequences labeled "…D"). Sequences harboring a variant TLR recognition motif (GTTG*C*GT) are labeled with a grey diamond. Note that most of those sequences are among the evolutionarily youngest HERV-K(HML-2) sequences, based on sequence divergence. Sequence designations follow a scheme "chr - start - end - chromosomal band". Sequence designations deviating from that scheme are polymorphic, non-reference HERV-K(HML-2) sequences. Genbank accession numbers and chromosomal bands or popular names are given for the latter.

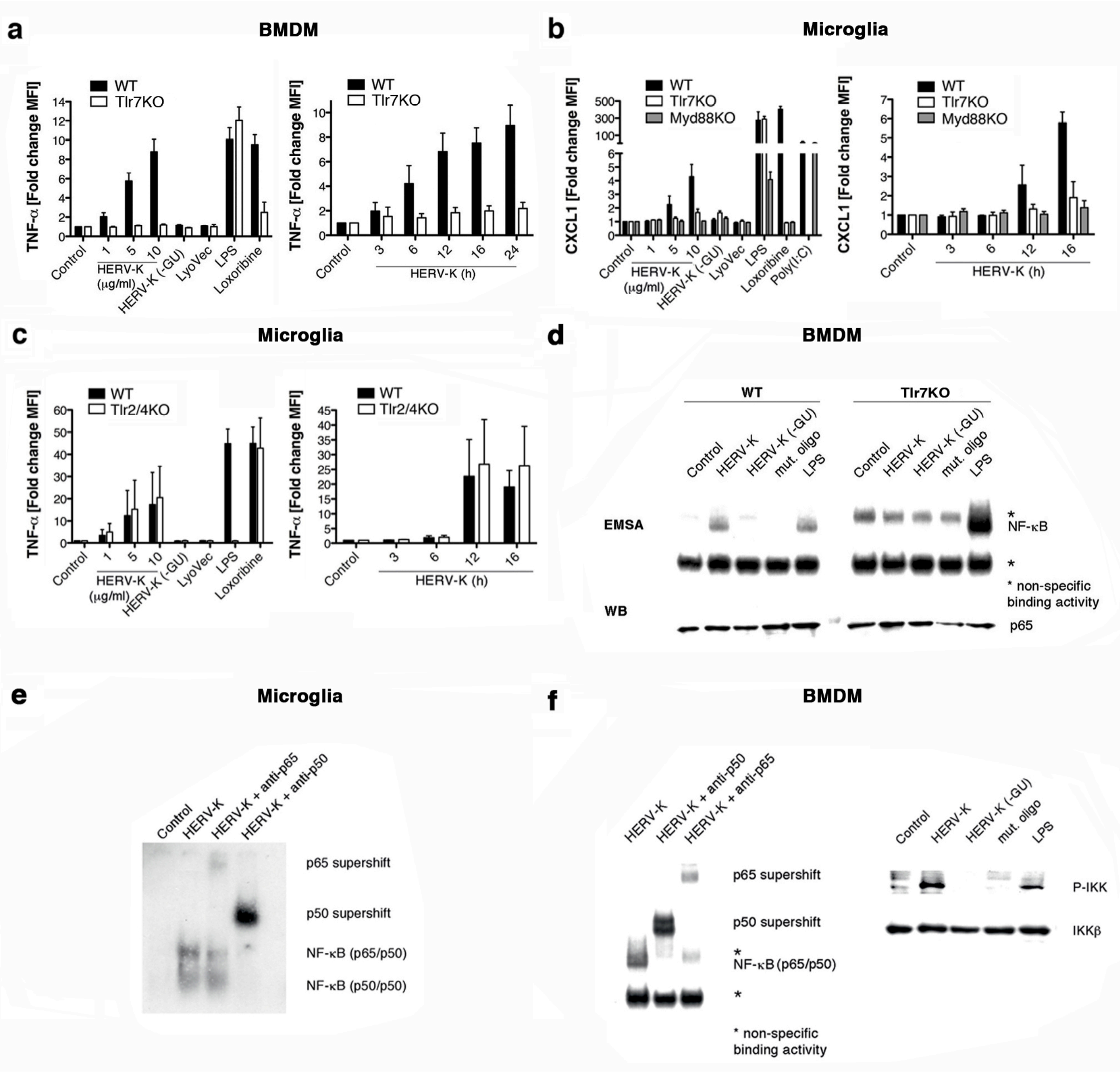


**Supplementary Figure 2. HERV-K RNA harboring a GU-rich sequence motif activates Tlr7 in microglia and macrophages, thereby inducing TNF-α and CXCL1 release through NF-κB signaling.** (**a**) Immortalized wild-type (WT) or Tlr7KO bone marrow-derived macrophages (BMDM), (**b**) microglia from C57BL/6J (WT), Tlr7KO, or Myd88KO mice, and (**c**) microglia from WT or Tlr2/4KO mice were incubated for 12 h with various doses of HERV-K(HML-2) (abbreviated as HERV-K in the following) oligoribonucleotide harboring a GUUGUGU motif within the *env* region (HERV-K, left) or with 5 μg/ml of

HERV-K for various durations (right). Untreated cells (control) and an oligoribonucleotide with a sequence located upstream of the GU-rich motif within the HERV-K *env* region (HERV-K (-GU), 10 µg/ml) served as a blank and negative control, respectively. LPS (100 ng/ml), loxoribine (1 mM), and poly:IC (100 ng/ml) were used as established ligands for TLR4, TLR7, and TLR3 activation, respectively. Subsequently, TNF-α (**a, c**) and CXCL1 (**b**) amounts in the culture supernatants were determined by immuno multiplex assay. Data are pooled from $n$ = 3-9 experiments. Results are presented as mean ± s.e.m. (**d**) BMDM were incubated for 2 h with 5 µg/ml HERV-K, 5 µg/ml HERV-K (-GU), or a mutant oligoribonucleotide, in which the GU content had been reduced. 1 µg/ml LPS served as a positive control. Subsequently, protein lysates were assayed for NF-κB activation by EMSA. Western blot using an antibody against p65 confirmed equal loading of probes. One representative experiment of at least three independent experiments is shown. **(e)** WT microglia and (**f**) BMDM were incubated with 5 µg/ml HERV-K, 5 µg/ml HERV-K (-GU), or mutant oligoribonucleotide for 2 h. 1 µg/ml LPS served as a positive control. Subsequently, cell lysates were assayed for NF-κB activation by EMSA. Western blots using antibodies against p65, p50, or p-IKK confirmed NF-κB activation. One representative experiment of at least three independent experiments is shown.

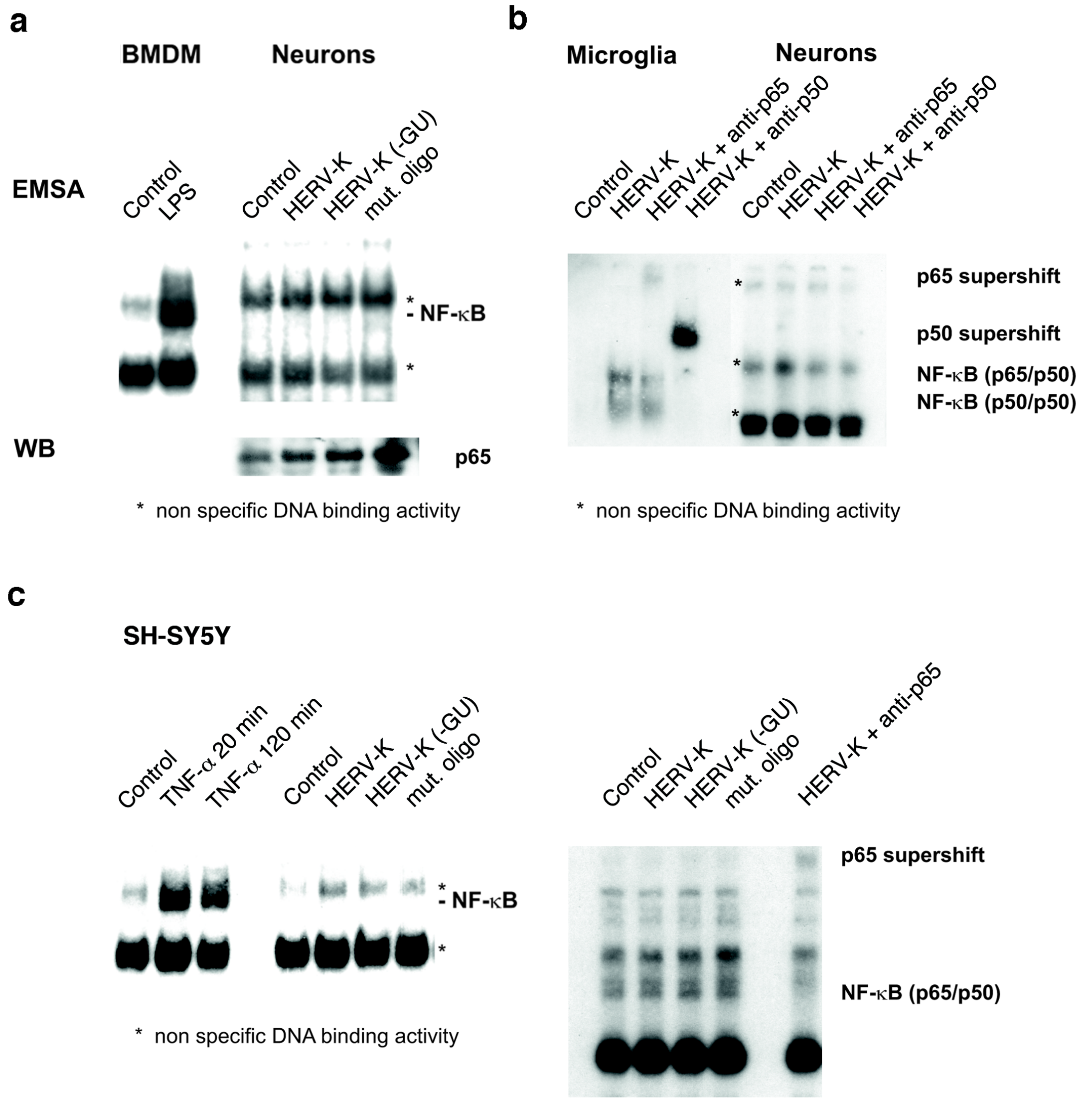


**Supplementary Figure 3. Stimulation of murine and human neurons with HERV-K RNA does not result in NF-κB activation.** Purified murine cortical neurons and (**a**) bone-marrow derived macrophages (BMDM) or (**b**) microglia, or (**c**) SH-SY5Y cells were incubated with 5 µg/ml HERV-K, 5 µg/ml HERV-K (-GU), or a mutant oligoribonucleotide, in which the GU content had been reduced, for 2 h. Untreated cells served as blank control, whereas 1 µg/ml LPS served as a positive control for BMDM activation. TNF-α served as a positive control for neuronal NF-κB activation. Subsequently, protein lysates were assayed for NF-κB activation by EMSA. Western blots using an antibody against p65 or EMSA signals derived from non-specific DNA binding activity served as loading control. Supershifts were determined by using antibodies against p50 and p65. One representative experiment of at least three independent experiments is shown.

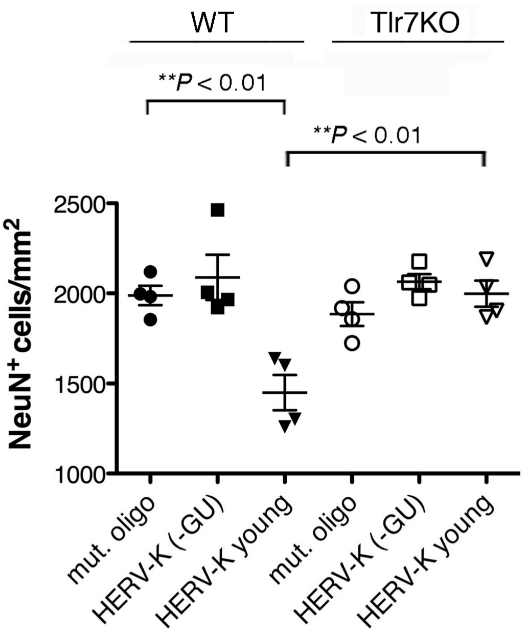


**Supplementary Figure 4. Intrathecal HERV-K RNA harboring the motif GUUG*C*GU induces neurodegeneration through Tlr7.** 10 µg HERV-K containing the sequence GUUG*C*GU (HERV-K young), 10 µg HERV-K (-GU) oligoribonucleotide, or 10 µg mutant oligoribonucleotide were injected intrathecally into C57BL/6J (WT, HERV-K young $n$ = 4; HERV-K (-GU) $n$ = 4; mutant oligoribonucleotide $n$ = 4) or Tlr7-deficient (Tlr7KO, HERV-K young $n$ = 4; HERV-K (-GU) $n$ = 4; mutant oligoribonucleotide $n$ = 4) mice. After 3 d, brain sections were immunostained with NeuN antibody. NeuN-positive cortical cells were quantified. Results are presented as mean ± s.e.m. A one-way ANOVA test yielded values of $P$ = 0.0003 over all groups. $P$ values for relevant groups were determined by one-way ANOVA with Bonferroni's post hoc test, as indicated.

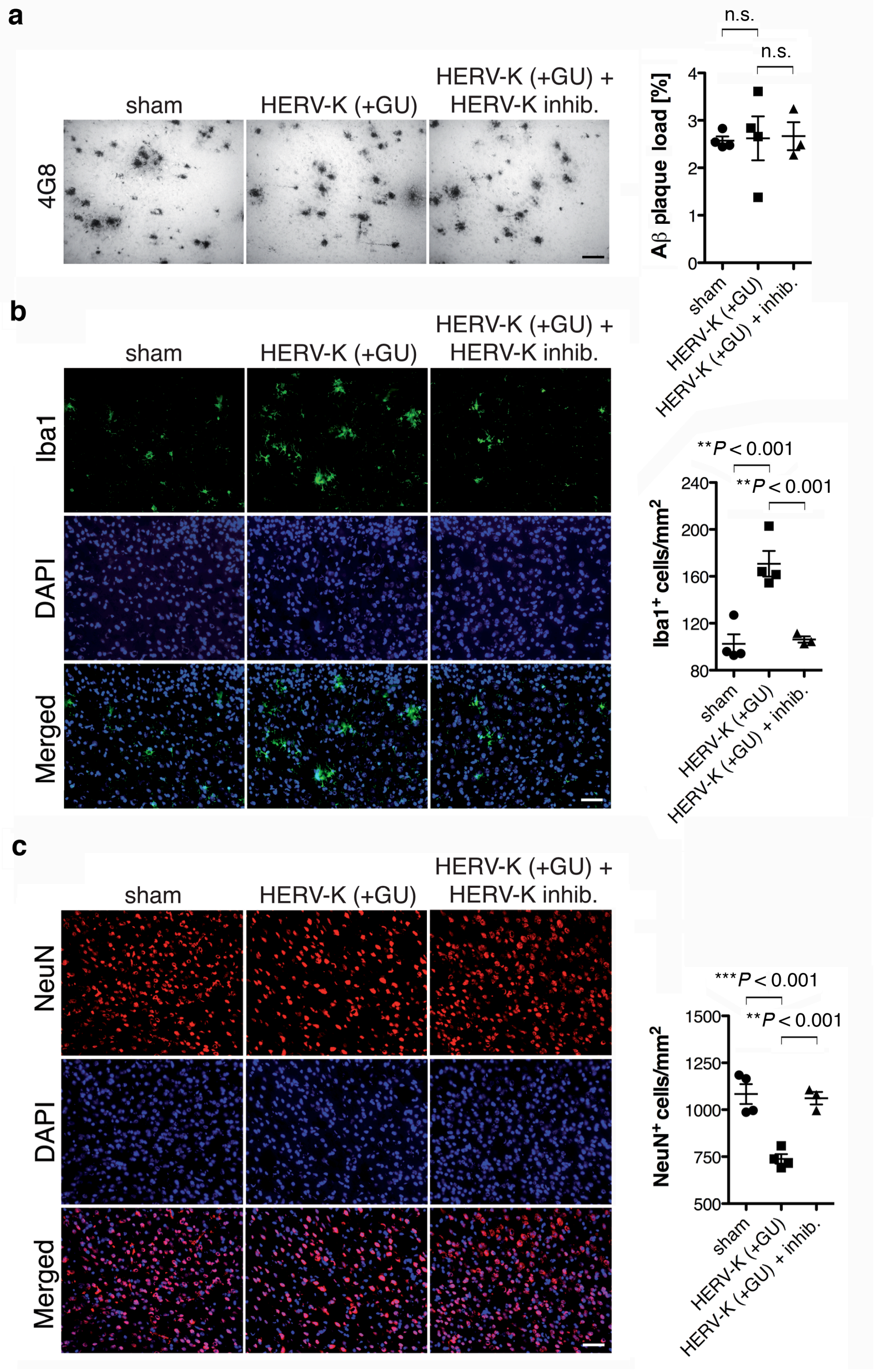


**Supplementary Figure 5. Extracellularly delivered HERV-K RNA results in increased numbers of microglia and loss of neurons in the cerebral cortex, but has no impact on**

**Aβ plaque burden in *APPPS1* mice.** 125 pmol of HERV-K inhibitor were injected intrathecally into 4-week-old *APPPS1* mice. After 16 h, mice were injected intrathecally with 10 μg of HERV-K (+GU). Sham-operated mice received PBS instead of oligoribonucleotides intrathecally. Subsequently, treatment was repeated every 4 weeks until the end of the experiment (120 d of age; sham, $n$ = 4; HERV-K (+GU), $n$ = 4; HERV-K (+GU) + HERV-K inhibitor, $n$ = 3). Brain sections were immunostained with 4G8 (**a**), Iba1 (**b**) and NeuN (**c**) antibodies. Scale bar, 50 μm. 4G8-positive Aβ plaques as well as NeuN- and Iba1-positive cells in the cortex were quantified. Results are presented as mean ± s.e.m. A one-way ANOVA test yielded values of $P$ = 0.9805 (**a**), $P$ = 0.0008 (**b**), and $P$ = 0.0004 (**c**) over all groups. $P$ values for relevant groups were determined by one-way ANOVA with Bonferroni's post hoc test, as indicated. n.s., not significant.

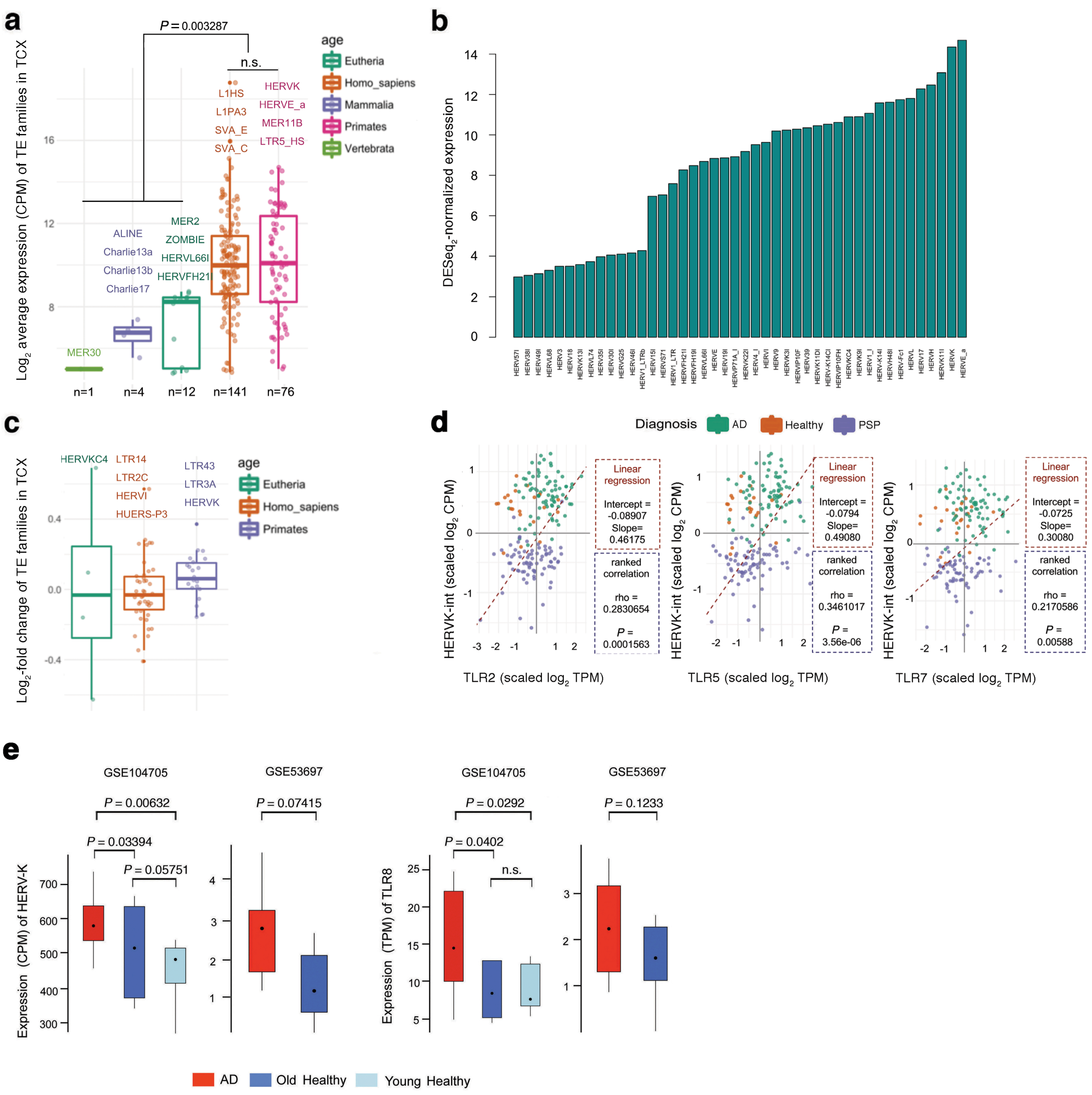


**Supplementary Figure 6. Specific expression of HERVs and TLRs in AD patients. (a)** Boxplots with jitters display scaled expression levels of various TE families in temporal cortex (TCX) samples (https://www.synapse.org/#!Synapse:syn4894912), based on clade annotations in Repbase. Dots indicate individual TE families of different phylogenetic ages, expressed as averages above the threshold ($\log_2$ CPM >2) across TCX samples. Numbers of expressed TE families, as well as their expression values, are significantly higher in the phylogenetically youngest categories (e.g. specifically expressed in primates/humans; ~200 out of ~750 families), compared to more ancient elements, shared by vertebrates/mammals/Eutheria) (17 out of ~400). Differential expression between human vs. primate-specific TEs was not significant. Note that most TEs expressed at the highest level were L1, SVA (non-LTR), HERVE_a and HERV-K (LTR). *P* values for relevant groups were determined by Wilcoxon test, as indicated. n.s. not significant. (**b**) Barplot displays the average expression of HERV families in the analysed TCX cohort. Only HERVs with $\log_2$ CPM >4 are plotted. (**c**) Boxplots with jitters display the level of differential expression of various HERVs based on clade annotations in Repbase. Dots indicate individual HERVs, expressed as average above threshold ($\log_2$ CPM >2 and $P$ <0.05, limma/Voom analysis) across TCX samples. (**d**) Correlation analysis of HERV-K expression and expression of various TLRs (TLR2, left; TLR5, middle; TLR7, right). Scatter plot shows scaled $\log_2$-transformed expression of HERV-K (CPM) and TLR (TPM) in TCX samples of AD, PSP and healthy ageing controls, as defined in (**a**). Linear regression analysis (dashed line) indicates correlation between HERV-K and TLR expression in AD samples. The rho value is obtained from pairwise-ranked correlation analysis. (**e**) Boxplots depicting distribution of HERV-K (left 2 graphs) and TLR8 (right 2 graphs) expression in brain samples from AD patients vs. healthy control individuals derived from previously published datasets: AD patients ($n$ = 12), young healthy ($n$ = 8) and aged healthy (Old healthy, $n$ = 10) individuals (lateral temporal lobe samples; GSE104705); patients with advanced AD ($n$ = 9) and healthy control individuals ($n$ = 8) (dorsolateral pre-frontal cortex samples, GSE53697). *P* values were determined by a two-tailed Student's *t*-test, which was further adjusted for multiple corrections, as indicated. n.s., not significant.

| Patient ID | Diagnosis | Gender | Age [years] | MMSE | ApoE | Total tau [pg·ml$^{-1}$][a] | Amyloid $\beta_{1-42}$ [pg·ml$^{-1}$][b] |
|---|---|---|---|---|---|---|---|
| C1 | Non-demented control | Male | 68 | 28 | 3/3 | 522 | 1220 |
| C2 | Non-demented control | Male | 64 | 29 | 3/3 | 243 | 806 |
| C3 | Non-demented control | Male | 59 | 28 | 3/3 | 239 | 947 |
| C4 | Non-demented control | Male | 74 | 29 | nd | 622 | 863 |
| C5 | Non-demented control | Female | 75 | 29 | 3/3 | 341 | 633 |
| C6 | Non-demented control | Female | 73 | 29 | nd | 107 | 576 |
| C7 | Non-demented control | Male | 76 | 28 | 3/4 | 338 | 342 |
| C8 | Non-demented control | Female | 58 | 28 | 3/3 | 208 | 529 |
| C9 | Non-demented control | Female | 66 | 27 | 3/3 | 209 | 541 |
| C10 | Non-demented control | Male | 63 | 30 | 3/3 | 427 | 869 |
| C11 | Non-demented control | Female | 77 | 29 | 3/4 | 555 | 631 |
| C12 | Non-demented control | Male | 68 | 30 | nd | 287 | 538 |
| C13 | Non-demented control | Female | 64 | 30 | 3/3 | 257 | 735 |
| C14 | Non-demented control | Female | 73 | 28 | 3/3 | 438 | 968 |
| C15 | Non-demented control | Female | 65 | 29 | 3/3 | 360 | 1172 |
| C16 | Non-demented control | Male | 73 | 27 | nd | 199 | 852 |
| C17 | Non-demented control | Male | 64 | 29 | nd | 190 | 755 |
| C18 | Non-demented control | Female | 78 | 29 | nd | 332 | 1139 |
| C19 | Non-demented control | Female | 63 | 30 | nd | 181 | 862 |
| C20 | Non-demented control | Female | 66 | 30 | nd | 200 | 669 |
| C21 | Non-demented control | Female | 70 | 29 | 3/4 | 307 | 1146 |
| C22 | Non-demented control | Male | 74 | 29 | 3/3 | 205 | 674 |
| C23 | Non-demented control | Male | 56 | 30 | 3/3 | 216 | 661 |
| C24 | Non-demented control | Female | 56 | 29 | 4/4 | 219 | 710 |
| C25 | Non-demented control | Male | 66 | 28 | nd | 174 | 728 |
| AD1 | Alzheimer's Disease | Male | 75 | 24 | 3/3 | 469 | 323 |
| AD2 | Alzheimer's Disease | Female | 66 | 18 | 3/4 | 820 | 250 |
| AD3 | Alzheimer's Disease | Male | 80 | 21 | 4/4 | 415 | 224 |
| AD4 | Alzheimer's Disease | Female | 73 | 23 | 3/4 | 672 | 309 |
| AD5 | Alzheimer's Disease | Male | 63 | 25 | 3/4 | 577 | 456 |
| AD6 | Alzheimer's Disease | Male | 78 | 17 | nd | 274 | 350 |
| AD7 | Alzheimer's Disease | Female | 76 | 21 | 3/3 | 1265 | 231 |
| AD8 | Alzheimer's Disease | Female | 76 | 25 | 2/4 | 466 | 335 |
| AD9 | Alzheimer's Disease | Male | 86 | 23 | 3/3 | 2279 | 476 |
| AD10 | Alzheimer's Disease | Female | 76 | 23 | 2/4 | 461 | 245 |
| AD11 | Alzheimer's Disease | Female | 74 | 14 | 2/4 | 677 | 366 |
| AD12 | Alzheimer's Disease | Male | 75 | 21 | 4/4 | 534 | 394 |
| AD13 | Alzheimer's Disease | Female | 75 | 22 | 3/4 | 1012 | 473 |
| AD14 | Alzheimer's Disease | Female | 80 | 21 | 3/4 | 962 | 289 |
| AD15 | Alzheimer's Disease | Male | 73 | 23 | nd | 654 | 202 |
| AD16 | Alzheimer's Disease | Female | 74 | 11 | 3/4 | 576 | 343 |
| AD17 | Alzheimer's Disease | Female | 63 | 7 | 4/4 | 1825 | 363 |
| AD18 | Alzheimer's Disease | Male | 65 | 12 | 3/4 | 951 | 424 |
| AD19 | Alzheimer's Disease | Male | 73 | 14 | 3/4 | 965 | 366 |
| AD20 | Alzheimer's Disease | Male | 76 | 19 | 3/3 | 509 | 388 |
| AD21 | Alzheimer's Disease | Female | 77 | 13 | 3/3 | 1799 | 483 |
| AD22 | Alzheimer's Disease | Male | 81 | 21 | 3/4 | 1446 | 392 |
| AD23 | Alzheimer's Disease | Female | 80 | 20 | 2/4 | 612 | 424 |
| AD24 | Alzheimer's Disease | Female | 70 | 25 | nd | 1119 | 471 |
| AD25 | Alzheimer's Disease | Male | 80 | 22 | 3/4 | 725 | 327 |

MMSE, Mini-Mental State Examination; nd, not determined; [a]295,1 ± 127,8 for Non-demented controls and 1052,7 ± 464,2 (mean ± s.d.) for patients diagnosed with Alzheimer's disease; [b]782,6 ± 219,5 for Non-demented controls and 398,1 ± 49,4 (mean ± s.d.) Alzheimer's disease patients.

**Supplementary Table 1. Demographic, neuropsychometric and biomarker data of control subjects (C) and patients with Alzheimer's disease (AD) selected for CSF RT-PCR analysis.**
